# Editing *cis*-elements of *OsPHO1;2* improved phosphate transport and yield in rice

**DOI:** 10.1101/2024.11.28.625858

**Authors:** Kanika Maurya, Balaji Mani, Bhagat Singh, Ujjwal Sirohi, Aime Jaskolowski, Sandeep Sharma, Harsha Vardhan Tatiparthi, Satendra Kumar Mangrauthia, Renu Pandey, Yves Poirier, Jitender Giri

**Author notes:** Author for correspondence: Jitender Giri, National Institute of Plant Genome Research, Aruna Asaf Ali Marg, New Delhi-110067, India. **Email ID**: Kanika Maurya Balaji Mani Bhagat Singh Ujjwal Sirohi Aime Jaskolowski Sandeep Sharma Harsha Vardhan Tatiparthi Satendra Kumar Mangrauthia Renu Pandey Yves Poirier Jitender Giri.

## Abstract

Increasing grain yield is the main aim of crop improvement, which is globally impacted by the low availability of soil phosphate (Pi). Overexpressing Pi transporters for increasing Pi uptake often result in Pi toxicity and growth retardation. Despite advancements in genetic engineering, targeting *cis-*regulatory motifs of Pi transporters remains underexplored for understanding plant mechanisms and improving Pi status. Here, we demonstrated that the excision of the transcription inhibitor motif from the promoter of Pi transporter, *OsPHO1;2,* enhanced its expression and increased root-to-shoot Pi transport, leading to improved grain yield. Using, *in-silico* and DNA-protein interaction studies, we show the role of OsWRKY6 transcription factor in the negative regulation of *OsPHO1;2* expression by binding to the *cis*-regulatory element (*W-box*) present in its promoter. *oswrky6* knockout lines displayed a higher *OsPHO1;2* expressions, and improved shoot Pi levels. Further, we engineered the *OsPHO1;2* promoter for precisely removing the W-box and enhancing *OsPHO1;2* expression. The phenotypic and physiological evaluation at the vegetative stage shows that *OsPHO1;2* promoter edited (*OsPHO1;2:PE*) lines have increased shoot length, plant biomass and more root-shoot Pi export under low and normal P conditions. Notably, the ^33^Pi uptake assay reveals that *OsPHO1;2:PE* lines exhibited enhanced root Pi uptake, supported by higher root-associated Pi transporters (*OsPHTs*) expression. An extensive agronomical assessment revealed that *OsPHO1;2:PE* lines acquire increased seed and panicle numbers, thereby increasing yield without affecting seed quality. Our findings offer valuable insight into the potential of promoter editing to raise plants with better Pi use and improved crop yield.

## Introduction

Phosphorus (P) is an essential macronutrient for growth, development, and yield, especially in crop plants. It plays a vital role in physiological and metabolic pathways like photosynthesis, respiration, and cellular signaling (Yang *et al*., 2024). Plants primarily acquire soluble inorganic orthophosphate (Pi) from soil. However, Pi in soils gets readily precipitated by other minerals (iron and aluminium) into non-soluble complexes, and microbes assimilate it into organic compounds (López-Arredondo *et al*., 2014). Due to the low availability of soil Pi, crops typically experience Pi deficiency, which hampers plant growth and productivity worldwide, making agriculture rely heavily upon the application of Pi fertilizers to attain high productivity (Jiao *et al*., 2012, Elser *et al*., 2017). However, due to the finite P reserves, high cost and environmental risks, like eutrophication, depending solely on Pi fertilizer is challenging and non-sustainable (MacDonald *et al*., 2011; Lun *et al*., 2018; Campos-Soriano *et al*., 2020). Therefore, it is necessary to understand the mechanisms behind plant responses to Pi starvation and utilize them for crop improvement targeted at efficient use of P.

Among various adaptive responses at the morphological, biochemical, and physiological levels, fine-tuning the expression of phosphate transporters in response to low Pi availability is critical for plants (Poirier *et al*., 2022). Root P acquisition from the rhizosphere is an energy-requiring process executed by the epidermal or cortical cells expressing H+/Pi co-transporters. In rice, at least thirteen members belonging to the PHT1 family have been identified expressing in root and involved in Pi acquisition (Mudge *et al*., 2002; Nussaume *et al*., 2011).

After absorption of Pi within the roots, Pi is translocated to the shoot by another class of Pi transporter expressed in xylem parenchyma cells named PHOSPHATE1 (PHO1), belonging to the SYG1/PHO81/XPR1-ERD1/XPR1/SYG1 (SPX-EXS) family (Poirier *et al*., 1991; Wang *et al*., 2004; Stefanovic *et al*., 2007). Of the three members of the rice PHO1 family (OsPHO1;1, OsPHO1;2, and OsPHO1;3), OsPHO1;2 is primarily involved in root-to-shoot Pi export (Secco *et al*., 2010). OsPHO1;2 is a plasma membrane-localized transporter that expresses in the root, young panicle, hulls, and developing seeds (Ma *et al*., 2021). *ospho1;2* mutants exhibit impaired Pi transfer, leading to excessive P accumulation in the roots, deficiency in the shoots, and underdeveloped grains (Secco *et al*., 2010; Che *et al*., 2020; Ko *et al*., 2024). Recently, Ma *et al*. (2024) demonstrated OsPHO1;2 functions in regulating leaf photosynthesis, and its mutation shows decreased electron transport activity and CO_2_ assimilation (Ma *et al*., 2024). The homologs of PHO1 in other crops, like SiPHO1 in tomatoes and members of CaPHO1 in chickpeas, play a major function in root-shoot Pi translocation (Zhao *et al*., 2019; Mani *et al*., 2024). PHO1 proteins harbour the SPX domain at the N-terminus, transmembrane α-helices in the middle, and the C-terminus EXS domain. Studies have shown that SPX proteins act as sensors for cellular Pi status by binding inositol pyrophosphate (PP-InsPs) molecules and regulating phosphate starvation response (PHR) (Nagpal *et al*., 2024; Zhao *et al*., 2024). Also, the EXS domain of AtPHO1;2 might aid in the uncoupling mechanism of Pi starvation and growth symptoms (Rouached *et al*., 2011; Wege *et al*., 2016). Such evidence highlights the additional role of PHO1 in plant P signaling, which is not yet well-understood.

Rice feeds more than half the world’s population, but almost 50 percent of rice-growing areas face Pi scarcity (Navea *et al*., 2024). Plants overexpressing PHTs or mutating PHT inhibitors can potentially increase the phosphate acquisition efficiency (PAE). However, increases in Pi uptake often cause Pi toxicity and reduced growth (Gu *et al*., 2016). For example, overexpressing *OsPHT1;2, OsPHT1;8* or *OsPHT1;9* increases Pi uptake and transport but also leads to Pi toxicity and reduces biomass and yield, especially under Pi sufficient conditions (Catarecha *et al*., 2007; Ai *et al*., 2009; Wang *et al*., 2014). Optimizing PHT1 expression may prevent Pi overaccumulation and enhance P starvation tolerance. Gene knockouts generated via CRISPR/Cas9 carry genetic variations leading to undesired pleiotropic effects. Editing *cis-* regulatory elements in the gene promoter is a promising tool for regulating gene expression and generating desired traits while avoiding undesired effects.

Details on WRKY domain transcription factors (TFs) mediated regulation of PHT transporters are emerging. These TFs contain the conserved domain WRKYGQK and bind to *W-box* or *W-box like cis*-element motif “TTGACC/T” in the promoter of various genes, including *PHTs*. WRKY TFs are involved in diverse processes influencing growth and plant signalling (Ulker and Somssich, 2004; Jiang *et al*., 2017). For instance, OsWRKY75 maintains tolerance to Pi starvation by regulating various OsPHT transporters (Devaiah *et al*., 2007). In Arabidopsis, *AtPHO1* is negatively regulated by AtWRKY6 and AtWRKY42. The double mutant *atwrky6atwrky42* showed increased expression of *AtPHO1* expression and a greater Pi transport to shoot (Chen *et al*., 2009; Su *et al*., 2015; Ye *et al*., 2018). Despite this, our current knowledge of the transcriptional regulation of rice *PHO1;2* remains elusive.

Here, we uncovered the molecular mechanism underlying the negative regulation of *OsPHO1;2* expression and presented an attractive approach for increasing its expression by modifying the *cis-*regulatory motif in its promoter. We report that the promoter of *OsPHO1;2* contains a W-box element recognized by the OsWRKY6 and suppresses *OsPHO1;2* expression and shoot Pi transfer. We applied CRISPR/Cas9-based promoter editing using two gRNAs to excise the *W-box* site for increasing *OsPHO1;2* expression while reducing the pleiotropic effects that could be associated with *oswrky6* knockouts. Our results further revealed that *W-box* removal increases the *OsPHO1;2* abundance, shoot Pi transfer, enhances Pi uptake, and overall plant P status without compromising seed quality. Moreover, *W-box* removal strategy increases panicle number and, ultimately, grain yield by up to 26%. These findings thus highlight the potential application of promoter editing strategy to improve crop productivity.

## Results

### OSWRKY6 binds to *OsPHO1;2* promoter and regulates its expression negatively

In Arabidopsis, *PHO1* is transcriptionally regulated by WRKYs (Chen *et al*., 2009; Ye *et al*., 2018). However, the regulation of *PHO1* in rice remains unclear. We found a potential *W-box* at –1280 bp in the promoter of rice phosphate transporter, *OsPHO1;2*, prompting an investigation into its transcriptional regulation.

In rice, there are 113 members of the WRKY TF family (Rushton *et al*., 2010). Gene co-expression analysis conducted using the public gene co-expression database RiceFREND (https://ricefrend.dna.affrc.go.jp/) to explore which WRKY TF potentially regulates *OsPHO1;2* expression, suggested that only one WRKY TF family member, OsWRKY6, co-expresses with *OsPHO1;2* (Table S1). In addition, OsWRKY6’s WRKY-domain shares 54% and 75.4% sequence identity and similarity, respectively (http://imed.med.ucm.es/cgi-bin/sias/) to AtWRKY6 protein, which has been studied for regulating *AtPHO1* expression (Chen *et al*., 2009; Ye *et al*., 2018). Moreover, phylogenetic analysis of the WRKY TFs of Arabidopsis and rice revealed that OsWRKY6 is one of the closest homologs to AtWRKY6 (Wu *et al*., 2005). Hence, all these data suggested that OsWRKY6 may have a role in regulating *OsPHO1;2* expression.

Transient expression of a eYFP-WRKY6 fusion in *N. bethamiana* demonstrated localization to the nucleus (Figure S1). To explore the expression pattern of *OsWRKY6,* we analysed its transcript levels in different developmental tissues. *OsWRKY6* expresses significantly higher in seeds, followed by roots and stem base (Figure 1a). This expression pattern was further validated by histological β-glucuronidase (GUS) analysis in different tissues of *OsWRKY6-promoter: GUS* reporter lines (Figure 1b). Interestingly, the transverse section of the roots showed the GUS expression specifically in stelar region coinciding with *OsPHO1;2* role in xylem loading (Figure 1b(ii)).

**Figure 1.**
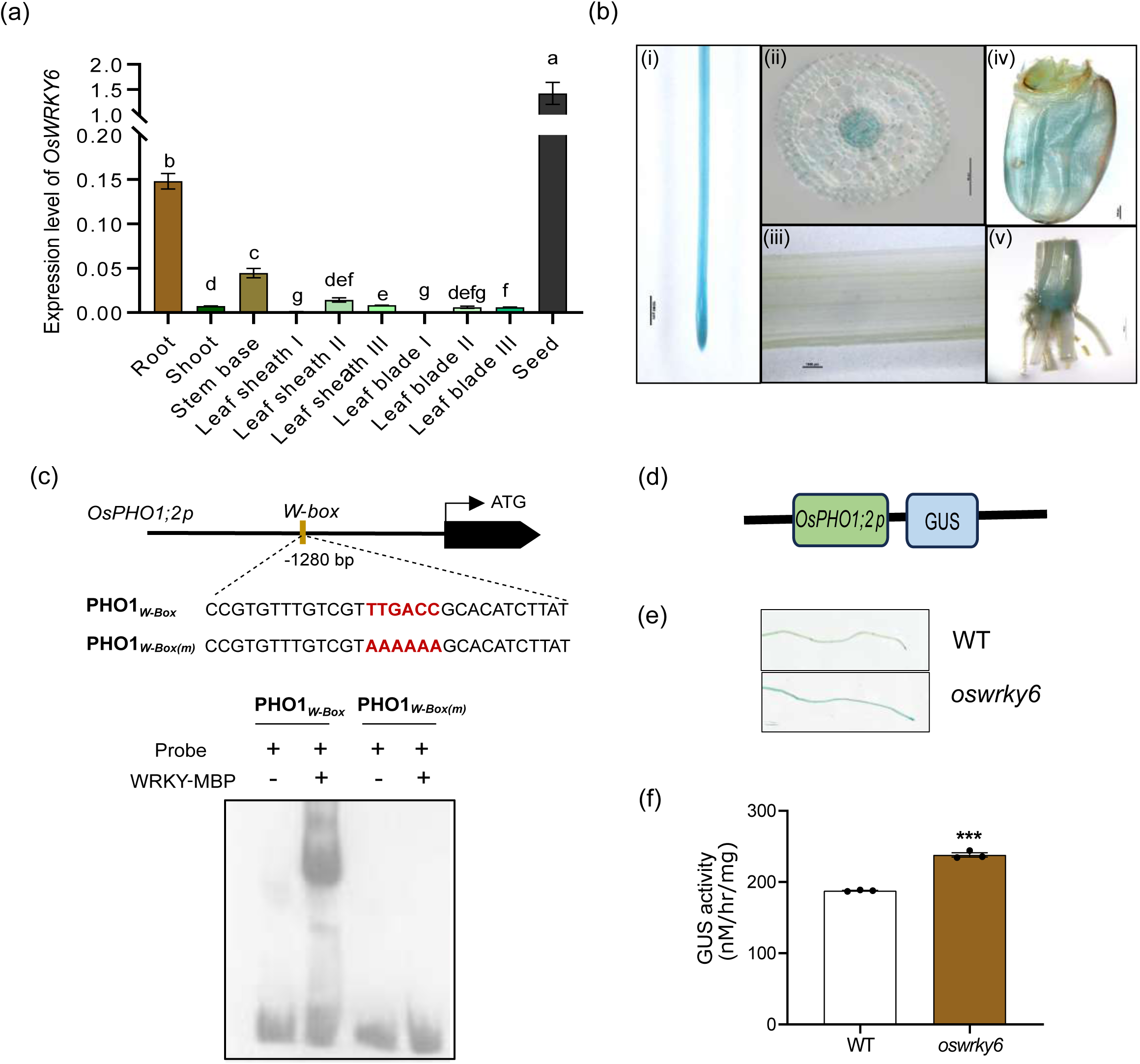
***OsWRKY6* co-expresses with *OsPHO1;2* and regulates its expression.** (a) The expression pattern of *OsWRKY6* in different rice tissues was determined by RT-qPCR. *Ubiquitin5* was used as an endogenous control. Data represent means ± SE (n = 3, each replicate contains a pool of 5 seedlings). Significant differences between different tissues are indicated with different letters (P < 0.05, one-way ANOVA) (b) Tissue-specific localization in ((i) root, (ii) transverse section of root, (iii) shoot, (iv) seed, (v) stem base) of *OsWRKY6* using GUS reporter lines driven by *OsWRKY6* promoter. (c) Electrophoretic mobility shift assay (EMSA) showing OsWRKY6 binding to *W-box* (PHO1*_W-Box_*) in *OsPHO1;2* promoter in vitro. The *W-box* mutated promoter sequence (PHO1*_W-Box(m)_*), in which the TTGACC mutated to AAAAAA, didn’t show the mobility shift with OsWRKY6. (d-f) Transient expression assay in rice seedlings showing repressor activity of OsWRKY6 on *OsPHO1;2* expression. (d) Reporter vector showing GUS (β-glucuronidase) gene driven by the promoter of the *OsPHO1;2*. (e) Images of 3-d-old rice seedlings (wild type (WT) and *oswrky6*) transformed with *OsPHO1;2p*:*GUS* construct followed by GUS staining. (f) Quantitative measurement of GUS activity in rice roots. Data represent means ± SE (n = 3, each replicate contains a pool of 5 seedlings). Each dot represents one biological replicate. Significant changes were determined using the Student’s *t*-test. *** indicates a significant difference from WT at P –value ≤ 0.001.

We next investigated whether OsWRKY6 can bind directly to *W-box* in *OsPHO1;2* promoter *(OsPHO1;2p)* using electrochemical mobility shift assay (EMSA). Our results showed that OsWRKY6 can bind to the *W-box* site in the *OsPHO1;2p*. However, mutation of *W-box* attenuates OsWRKY6 binding to *OsPHO1p* (Figure 1c). To understand how OsWRKY6 regulates *OsPHO1;2* expression, we transiently expressed *OsPHO1;2p:GUS* construct in the rice roots of wild type (WT) and *oswrky6* knockout. As shown in Figure 1d, *oswrky6* knockout roots resulted in a strong GUS signal compared to WT, as visualized by GUS staining and further from the quantification of GUS activity (Figure 1d-f).

These findings suggest that OsWRKY6 negatively regulates *OsPHO1;2* expressions by directly interacting with the *W-box* present in the *OsPHO1;2* promoter.

### Knocking out *OsWRKY6* increases *OsPHO1;2* expression in roots

To better understand the role of OsWRKY6 in *OsPHO1;2* regulation, we targeted the native *OsWRKY6* gene using CRISPR/Cas9 tool and generated the *oswrky6* mutant lines. We selected two lines based on the sequencing results of the target site, having 1bp deletion leading to null mutation (Figure S2). The transcript levels of *OsWRKY6* in these mutants were reduced compared to the WT (Figure S3). Next, to examine the effect of OsWRKY6 knockout on the regulation of the *OsPHO1;2* expression and to validate our *in vitro* data, we examined the *OsPHO1;2* expression in the roots of these *oswrky6* mutants. Remarkably, the *oswrky6* mutants exhibited a higher *OsPHO1;2* transcript levels in roots compared to the WT (Figure 2a). Consequently, the improved *OsPHO1;2* expression was associated with a higher Pi accumulation in the shoots of *oswrky6* mutants than WT (Figure 2b). *oswrky6* knockouts showed up to ∼27% and ∼21% increased shoot Pi under low and normal P conditions, respectively. However, phenotypic analysis revealed that *oswrky6* mutants grew like WT and didn’t show any significant difference in the shoot length and plant biomass under different Pi conditions (Figure S4).

**Figure 2.**
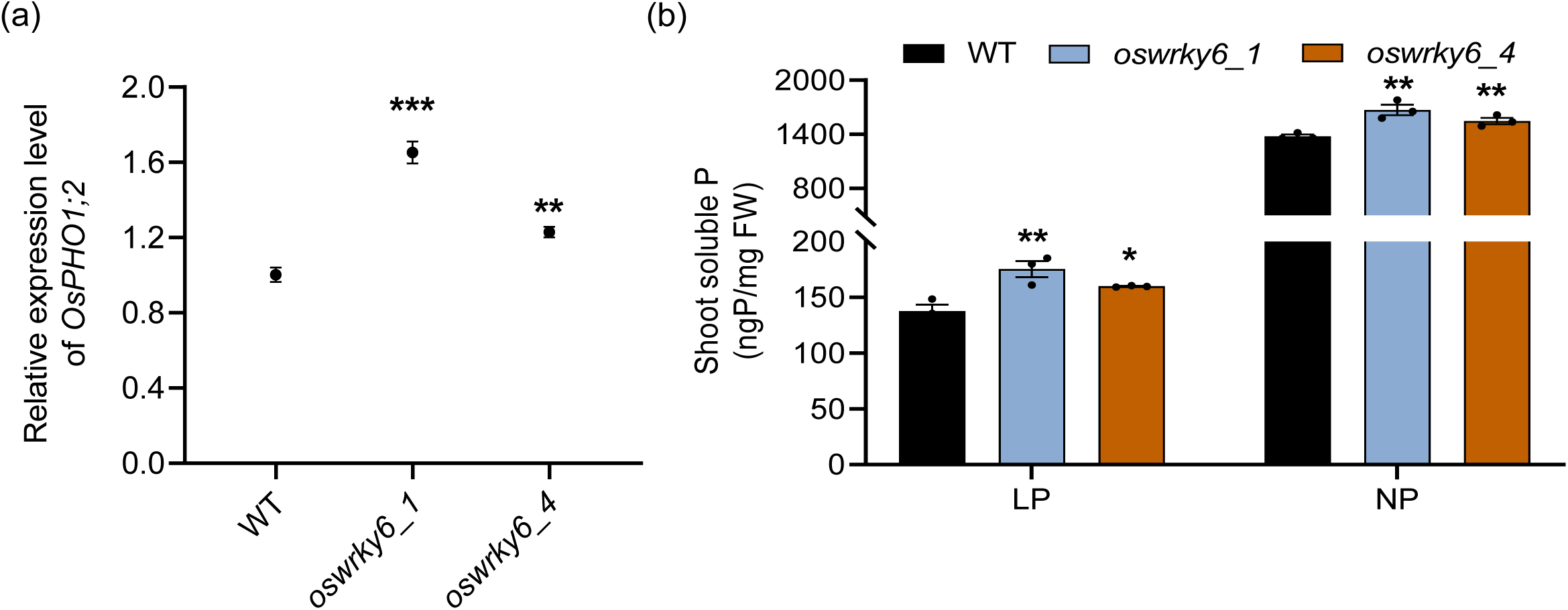
***oswrky6* knockout lines show enhanced *OsPHO1;2* expression with improved shoot Pi concentration**. (a) Relative expression level of *OsPHO1;2* in root tissue of 9-days-old seedlings of wild type (WT) and *oswrky6* mutants. Expression level was calculated relative to WT. *Ubiquitin5* was used as an endogenous control. Data represent means ± SE (n=4, each replicate contains a pool of 5 seedlings). (b) Soluble P estimation in shoots of 21-days-old seedlings of WT and *oswrky6* mutants grown under low (10 µM NaH_2_PO_4,_ LP) and normal P (320 µM NaH_2_PO_4,_ NP) conditions. Data represent means ± SE (n = 3, each replicate contains a pool of 5 seedlings). Each dot represents one biological replicate. Significant changes were determined using the Student’s *t*-test. *, ** and *** indicate significant difference from control (WT) at P-value ≤0.05, ≤0.01 and ≤0.001, respectively.

These results indicate that OsWRKY6 reduced Pi transfer to shoot by negatively regulating *OsPHO1;2* activity and mutation of OsWRKY6 improves Pi transfer to shoot with intriguingly no discernible changes in plant growth.

### Precise removal of OsWRKY6 binding site from *OsPHO1;2*’s promoter (*OsPHO1;2p*) contributes to enhanced *OsPHO1;2* expression and improves shoot Pi

Based on the above results, shoot P levels can be enhanced by overexpressing *OsPHO1;2* or suppressing *OsWRKY6*. Targeted promoter editing is an interesting alternative to fine-tune the gene expression and create new traits without generating loss of function mutant or introducing transgene. In this regard, we aimed to increase the expression of *OsPHO1;2* by removing the *W-box* from its promoter using CRISPR/Cas9.

Two closest gRNAs flanking the *W-box* region in *OsPHO1;2p* were designed, and stable transgenic rice plants were generated (Figure 3a). The target region was examined for editing using fragment-specific PCR, and we obtained transgenic plants containing a deletion of ∼30 bp in *OsPHO1;2* promoter (Figure S5). We selected two independent lines in T2 generation showing the desired biallelic deletion and performed DNA sequencing. Sequencing results revealed two types of deletions in the transgenic lines: 33 bp (*OsPHO1;2:PE8*) and 35 bp (*OsPHO1;2:PE5*). Both deletions occur in the *W-box* region of the *OsPHO1;2p*. (Figure 3b). The progenies from these lines were screened, and CRISPR/Cas9 free lines were selected (Figure S6). Further, we also studied potential off-target mutations using the CRISPR GE tool (http://skl.scau.edu.cn/) and detected 13 and 12 putative off-targets for gRNA1 and gRNA2, respectively. Most off-targets were found in intergenic and intronic regions with > 4 mismatches in the seed region (Table S2). The top five off-targets showing the highest off-target scores were selected and screened. Upon screening a pool of 10-15 plants, much lower off-targets score was obtained for both *OsPHO1;2:PE* lines (Table S3). Also, the distribution of off-targets mutation occurs in non-coding regions, suggesting high specificity of designed gRNAs for *W-box* excision.

**Figure 3.**
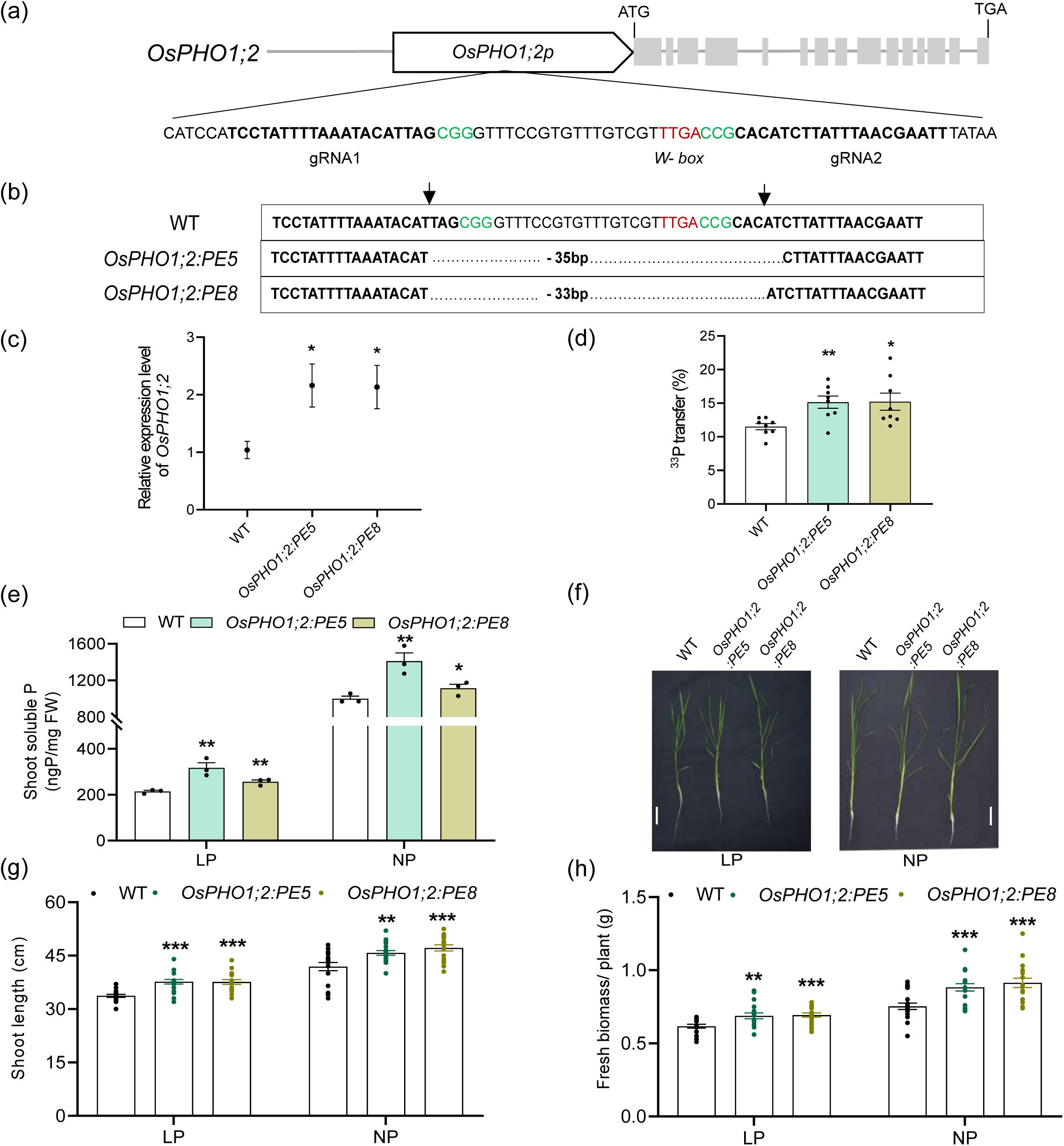
**Promoter editing of *OsPHO1;2* enhances *OsPHO1;2* expression with improved shoot Pi transport and plant growth**. (a) Schematic illustration of *OsPHO1;2* gene and its promoter (*OsPHO1;2p*). The introns and exons are represented by grey-color line and rectangle boxes, respectively. The translation initiation codon (ATG) and termination codon (TGA) are shown. The W-box sequence (TTGACC) in *OsPHO1;2p* is represented by a red color nucleotide sequence. The gRNA sequences (gRNA1 and gRNA2) are underlined and bold. (b) Sanger sequencing analysis of *OsPHO1;2:PE* plants. The CRISPR/Cas9 mediated deletion of *OsPHO1;2p* sequence are indicated in base pairs. gRNA sequences are in bold, nucleotides marked in green represent PAM; arrow denotes putative Cas9 cut site (3bp upstream to PAM); *W-box* is represented by red colored and underlined sequence. For Sanger sequencing, the DNA fragment spanning the target sites was amplified using primers spanning gRNAs, followed by sequencing. (c) The expression level of *OsPHO1;2* in root tissue of 9-days-old seedlings of wild type (WT) and *OsPHO1;2:PE* lines. Relative expression was calculated relative to WT. *Ubiquitin5* was used as an endogenous control. Data represent means ± SE (n= 4, each replicate contains a pool of 5 seedlings). (d) ^33^P transfer rate from root to shoot in 9-d-old seedlings of WT and *OsPHO1;2:PE* lines. Data represent means ±SE (n = 8). (e) Soluble P estimation in shoot tissues of 21-days-old seedlings of WT and *OsPHO1;2:PE* lines grown under low (10 µM NaH_2_PO_4,_ LP) and normal P (320 µM NaH_2_PO_4,_ NP) conditions. Data represent means ± SE (n= 3). (f) Phenotypic analysis, (g) shoot length and (h) plant biomass of 21-days-old seedlings of WT and *OsPHO1;2:PE* lines grown under LP and NP conditions. Data represent means ± SE (n=15-20). Each dot represents one biological replicate. Scale bar: 5cm. Significant changes were determined by the Student’s *t*-test. *, ** and *** indicate significant difference from control (WT) at P-value ≤0.05, ≤0.01 and ≤0.001, respectively.

Next, we studied the effect of *OsPHO1;2p* editing on *OsPHO1;2* gene expression by performing RT-qPCR in the root tissues of *OsPHO1;2:PE* lines (Figure 3c). Our results showed that *OsPHO1;2* expression is elevated significantly in both *OsPHO1;2:PE* lines, suggesting that removal of *W-box* positively regulates *OsPHO1;2* expression. Further, the ^33^P transfer rate was examined in the *OsPHO1;2:PE* lines as OsPHO1;2 in rice is a major Pi exporter for root-shoot translocation (Secco *et al*., 2010). We obtained a 32% increase in root-shoot ^33^P transfer rate in *OsPHO1;2:PE* lines compared to the WT (Figure 3d). Additionally, increased expression of *OsPHO1;2* resulted in higher Pi levels in shoot tissue, with a substantial increase of up to 48% and 41% in shoots of *OsPHO1;2:PE* lines under low and normal P conditions, respectively (Figure 3e). These findings were further accompanied by a significantly greater total shoot P level in the *OsPHO1;2:PE* lines (Figure S7a).

Subsequently, we monitored the growth performance of *OsPHO1;2:PE* lines at the vegetative stage under different P concentrations (Figure 3f). Compared to WT, *OsPHO1;2:PE* lines showed >11.5% and >9% increment in shoot length under low and normal P conditions, respectively (Figure 3g). A similar trend was obtained in the case of plant biomass and shoot dry weight where *OsPHO1;2:PE* lines’ show increased up to 21% compared to WT (Figure 3h, S7b).

Therefore, we concluded that at the early vegetative stage, removal of the OsWRKY6 binding site, i.e., *W-box* (TTGACC/T), elevates *OsPHO1;2* expression, causing more Pi transfer to shoot and improved plant growth.

### *OsPHO1;2:PE* lines show higher root P accumulation with induction in root associated PHTs expression

Interestingly, when we looked at the root phenotype, *OsPHO1;2:PE* lines showed shorter roots in comparison to WT (Figure 4a). On the contrary, the root number in *OsPHO1;2:PE* lines increased by about 22-55% under both low and normal P conditions (Figure 4b). This made us examine Pi status of roots. Compared to WT, roots of *OsPHO1;2:PE* lines accumulate >40% and >34% high Pi under low and normal P conditions, respectively (Figure 4c). Similarly, we observed significantly more total P in the roots of *OsPHO1;2:PE* lines (Figure S8a). Further, we witnessed an enhanced ^33^P uptake up to 60% by the roots of *OsPHO1;2:PE* lines compared to WT, suggesting more Pi in root is related to higher Pi uptake mediated by root Pi transporters (Figure 4d). To validate this possibility, we studied the expression of root-associated *OsPHTs* in *OsPHO1;2:PE* lines and detected a significant increase in the expression of *OsPHT1;1, OsPHT1;4, OsPHT1;8* and *OsPHT1;9* (Figure 4e-i). Interestingly, a significant rise in the expression of the low-affinity Pi transporter, *OsPHT1;2* was noted. With this data, we calculated root efficiency in *OsPHO1;2:PE* lines, representing the total amount of P accumulated within the plant providing the same root surface area. In our observation, root efficiency is improved in *OsPHO1;2:PE* lines. Thus, enhanced *OsPHO1;2* expression resulted in high expression of root *OsPHTs*, leading to elevated P levels in roots in *W-box* edited lines, indicating better root efficiency in *OsPHO1;2:PE* lines. (Figure S8b).

**Figure 4.**
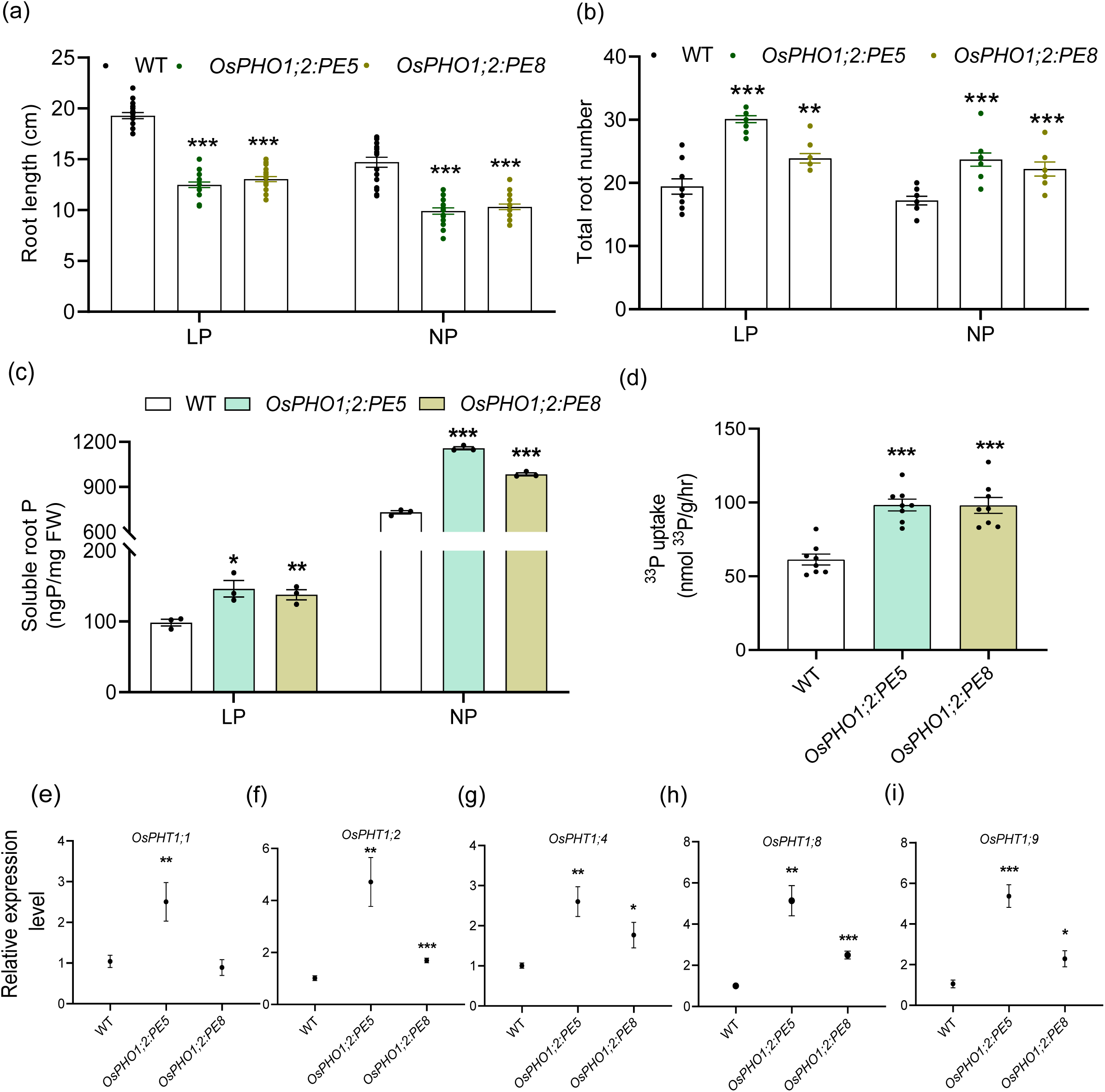
Promoter editing of *OsPHO1;2* increased Pi uptake. (a) Root length (n=15-20) and (b) total number of roots (n=8-10) measurement in 21-days-old seedlings of wild type (WT) and *OsPHO1;2:PE* lines grown under low (10 µM NaH_2_PO_4,_ LP) and normal P (320 µM NaH_2_PO_4,_ NP) conditions. (c) Root soluble P in 21-day-old seedlings of WT and *OsPHO1;2:PE* lines grown under LP and NP conditions. Data represent means ± SE (n= 3, each replicate contains a pool of 5 seedlings). (d) ^33^P uptake rate in 9-days-old seedlings of WT and *OsPHO1;2:PE* lines. ^33^P uptake rate represents the amount of ^33^P absorbed by the roots from the external solution per mg of root fresh weight. Data represent means ±SE (n=8). (e-i) Relative expression of phosphate transporters (*OsPHTs*); (e) *OsPHT1;1,* (f) *OsPHT1;2,* (g) *OsPHT1;4,* (h) *OsPHT1;8* and (i) *OsPHT1;9* in root tissue of 9 days-old seedlings of WT and *OsPHO1;2:PE* lines. Expression was calculated relative to WT. *Ubiquitin5* was used as an endogenous control. Data represent means ± SE (n= 4, each replicate contains a pool of 5 seedlings). Each dot represents one biological replicate. Significant changes were determined by the Student’s *t*-test. *, ** and *** indicate significant differences from control (WT) at P-value ≤0.05, ≤0.01 and ≤0.001, respectively.

### *W-box* removal of *OsPHO1;2* promoter improves panicle number and plant yield

Next, we evaluated the growth performance of *OsPHO1;2:PE* lines in soil-filled pots with normal P supply. To this end, we phenotyped the *OsPHO1;2:PE* lines in the year 2023 and 2024. In the 2023 season, the plant height, leaf length, leaf width flag leaf length, flag leaf width, and panicle length of *OsPHO1;2:PE* lines don’t show any significant difference compared to WT (Figure 5a, b, d, S9a-d). However, reproductive traits like tiller number, panicle number per plant in *OsPHO1;2:PE* lines were significantly increased by 26-45% compared to WT (Figure 5c, e). A closer examination of panicles revealed that the branch number per panicle and branch length had no change, whereas few panicle branches show a marginal increase in seeds/ branch in *OsPHO1;2:PE* lines (Figure S9e-g). Remarkably, the number of filled seeds/panicle was improved in *OsPHO1;2:PE* lines compared to WT, culminating in a higher seed yield (Figure 5f). A similar trend of a greater number of tillers, panicles, and yield was observed in the 2024 season (Figure S10). Also, we looked at the performance of *OsPHO1;2:PE* lines in soil fertilized with less phosphorus. Compared to WT, plant height, flag leaf width, and panicle length were reduced in *OsPHO1,2:PE* lines; however, these plants displayed a greater number of panicles and improved seed yield (Figure S11).

**Figure 5.**
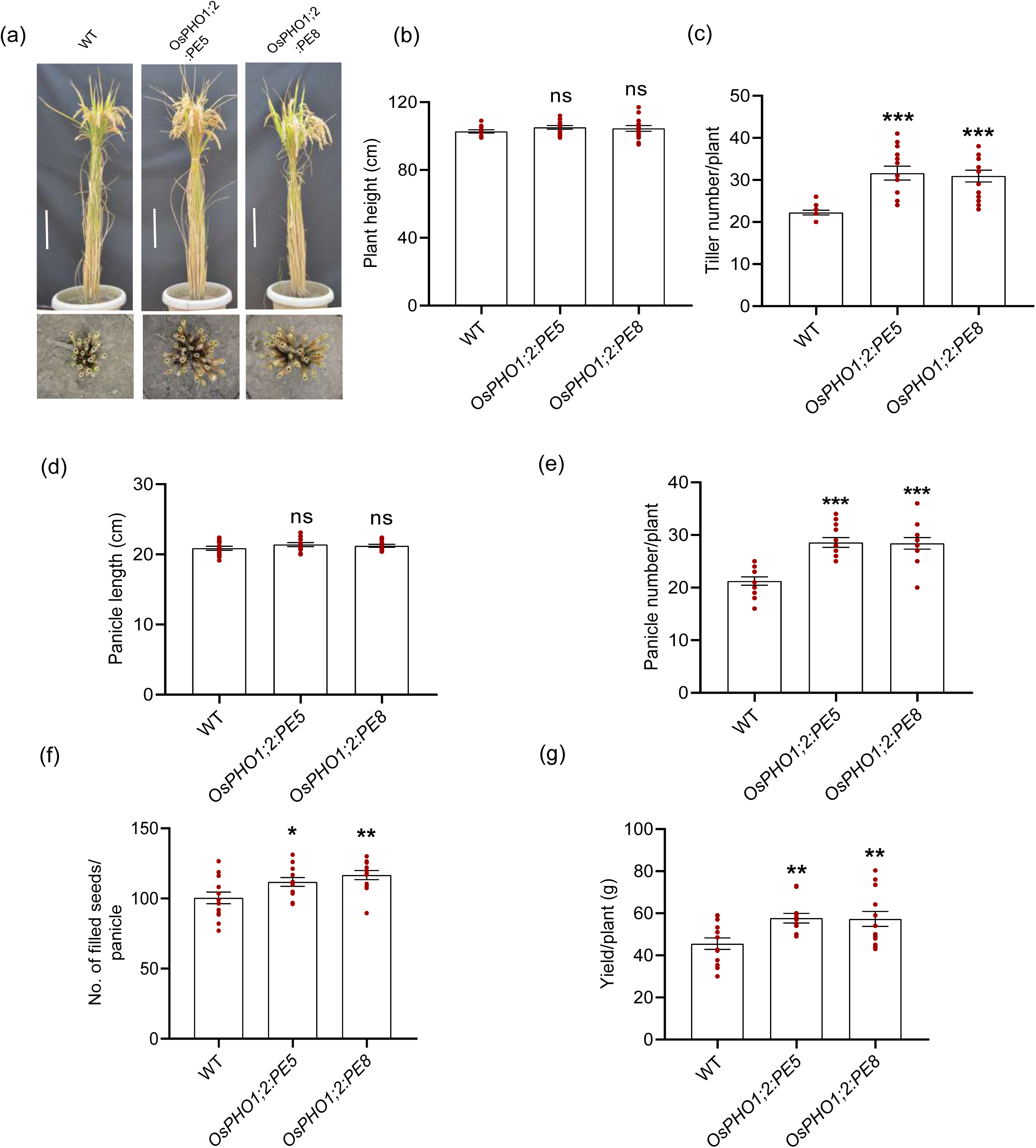
**Promoter editing of *OsPHO1;2* improves yield-related traits in rice (Year 2023)**. Agronomic performance of WT and *OsPHO1;2:PE* lines at mature stage; (a) Gross morphology and tillers of WT and *OsPHO1;2:PE* lines at mature stage, (b) plant height, (c) tiller number, (d) panicle length, (e) panicle number/ plant, (f) number of filled seeds/ panicle and (g) yield/ plant. Data represent means ± SE (n = 12-15). Each dot represents one biological replicate. Significant changes were determined by the Student’s *t*-test. ns indicate no significant difference. *, ** and *** indicate significant differences from control (WT) at P-value ≤0.05, ≤0.01 and ≤0.001, respectively. Scale bar: 20 cm.

Hence, our results showed that removing the *W-box* from the *OsPHO1;2* promoter increases panicle number, leading to higher grain yield in *OsPHO1;2:PE* lines under both low and normal P supply.

### *OsPHO1;2:PE* seeds have reduced P content with no compromise in quality

Apart from panicle number, seed number, and total plant yield, grain quality is a critical factor in rice for human consumption (Hori and Sun, 2022; Zhou *et al*., 2020). Since *OsPHO1;2:PE* lines showed higher yield because of increased panicle number per plant, we examined seed quality using parameters like seed size, length, and width in WT and *OsPHO1;2:PE* lines. Our results showed *OsPHO1;2:PE* seed dimensions were comparable to WT, and no significant variation was observed (Figure 6a-d). The primary component of rice endosperm is starch granules, which act as carbohydrate storage and ultimately determine the cooking quality (Yu *et al*., 2009; Zhu *et al*., 2023). Rice seeds were cut in half and observed under the scanning electron microscope to observe starch granule morphology. Most of the starch granules were polygonal in shape and tightly packed in seeds of both WT and *OsPHO1;2:PE* lines with no visible differences (Figure 6e). This result was further supported by the iodine staining of seeds and the quantification of starch and amylose content (Figure S12 and 6f).

**Figure 6.**
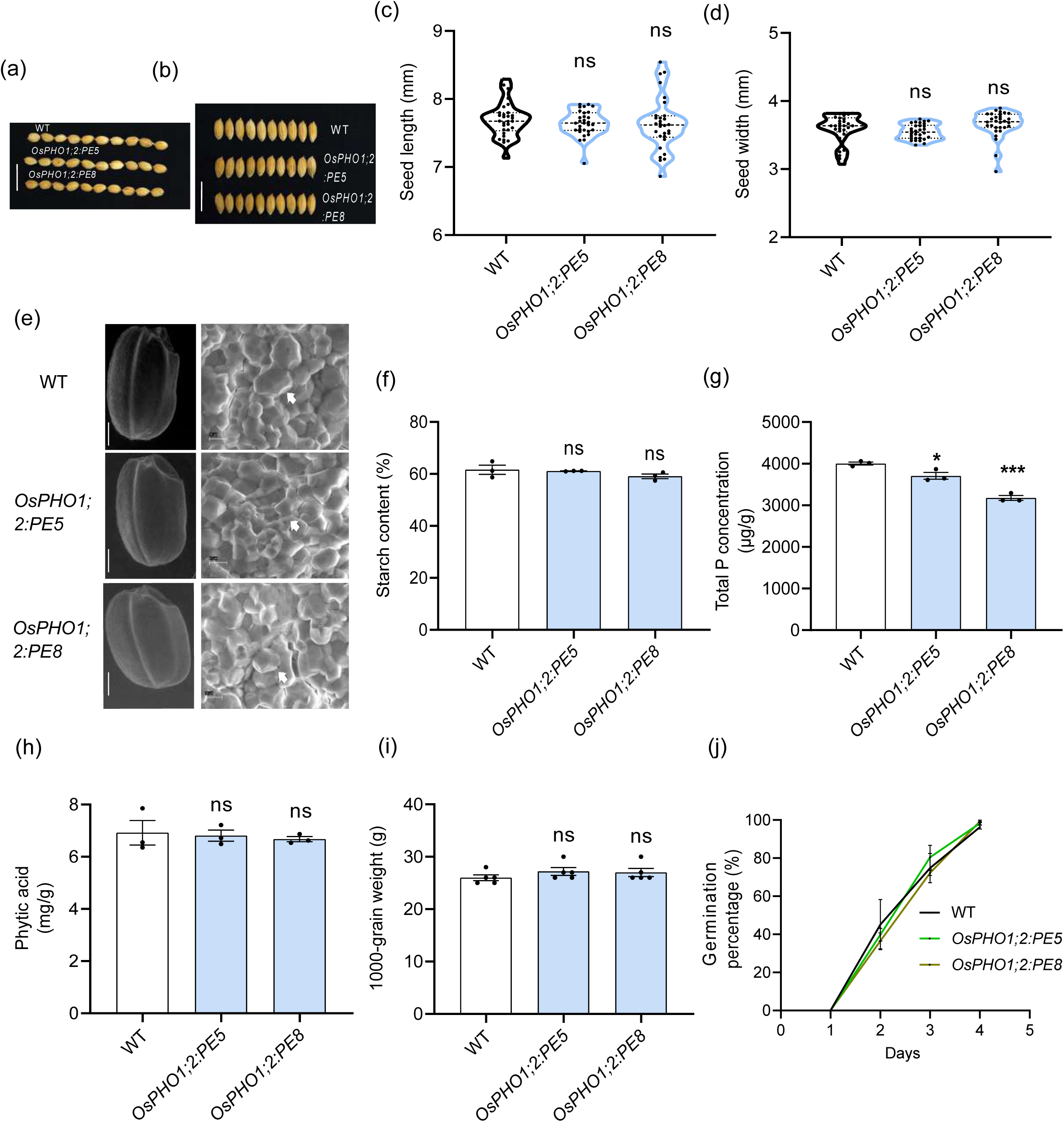
***OsPHO1;2:PE* lines show no penalty in seed quality**. (a) Seed length and (b) seed width phenotype of mature seeds of wild type (WT) and *OsPHO1;2:PE* lines. Scale bar:1cm. (c) Seed length and (d) seed width measurement of mature seeds of WT and *OsPHO1;2:PE* lines. (n=30 seeds). (e) Scanning electron microscopic images of whole seed (scale bar: 30 mm) and middle cross-sections (scale bar: 10 µm) of brown seeds of WT and *OsPHO1;2:PE* lines. Arrow indicates packaging of starch granules. Measurement of (f) starch content, (g) total P concentration and (h) phytic acid in brown seeds of WT and *OsPHO1;2:PE* lines. Data represent means ± SE (n=3). (i) 1000 grain weight (n=5) and (j) Germination percentage of WT and *OsPHO1;2:PE* lines (n=3, 30 plants in each replicate). Data represent means ± SE. Each dot represents one biological replicate. Significant changes were determined by the Student’s *t*-test. ns indicate no significant difference. * and *** indicate significant difference from control (WT) at P-value ≤0.05 and ≤0.001, respectively.

Recent reports have shown that the expression of *OsPHO1;2* influences seed development through a maternal effect by regulating Pi content in the endosperm, resulting in *ospho1;2* mutants having low starch content and shrunken seeds (Ko *et al*., 2024; Ma *et al*., 2021). We observed that *OsPHO1;2:PE* lines accumulate marginally lesser total seed P content than WT (Figure 6g). Also, the grain physiological phosphate use efficiency (PPUE) is increased in *OsPHO1;2:PE* lines, hinting that seed P content of *OsPHO1;2:PE* lines utilized in producing a greater number of seeds and eventually higher yield (Figure S13). Additionally, since the Pi and Fe show fine-tuning in regulating the plant growth (Lay-Pruitt *et al*., 2022), we profiled Fe content in seed where we observed such change in seed Pi concentration doesn’t have any impact on Fe content (Figure S14). Phytic acid is the principal storage P form in seeds and antinutrients for human consumption (Bloot et al., 2023; Nissar et al., 2017). Because of its chelating properties, the high phytic acid in seeds results in indigestion and other health-related issues. Thus, we looked at the phytic acid content of *OsPHO1;2:PE* lines, where we observed no discernible variation in the amount of phytic acid compared to WT (Figure 6h). Also, no difference was observed in 1000-seed weight and germination percentage in *OsPHO1;2:PE* lines compared to WT seeds (Figure 6i, j).

Collectively, we show that in *OsPHO1;2PE* lines produce completely normal seeds in terms of size and quality. Also, these promoter-edited lines do not display any reproductive defects.

## Discussion

Over the last century, grain production has heavily relied on phosphate fertilizers, boosting global crop yields. By 2050, as the world population approaches 10 billion, P fertilizer demand will exceed 22-27 Tg P yr^−1^. Agricultural P use efficiency (PUE) must rise to 80% to meet food demands (Zou *et al*., 2022). Rice exhibits low PUE among cereals, necessitating more fertilizer to combat hunger in countries like India (Zou *et al*., 2022). As a major P fertilizer importer, India needs to enhance its PUE through changes in agricultural practices (Langhans *et al*., 2022). Furthermore, global P fertilizer overapplication, exceeding optimal plant needs by 30-40%, causes nutrient runoff, leading to algal blooms and hypoxia (Lun *et al*., 2018; McDowell *et al*., 2024). Thus, effective crop production requires better fertilizer management and a deeper understanding of plant phosphate transport. Gene editing presents a groundbreaking method for advancing plant biology and addressing various agricultural issues (Ahmar *et al*., 2020). Our study utilized CRISPR/Cas9 to modify the cis-regulatory region of the phosphate transporter OsPHO1;2, enhancing P transport and yield under varying P conditions.

WRKY TFs are key players in plant signaling pathways linked with biotic and abiotic stresses (Wang *et al*., 2023; Wani *et al*., 2021; Zhang *et al*., 2021). Our initial sequence analysis showed that OsWRKY6 shares a high sequence similarity to the *AtPHO1* inhibitor, AtWRKY6, and co-expresses with *OsPHO1;2* in roots (Figure 1). Interestingly, by binding to the *W-box* site in the *OsPHO1;2* promoter, *OsWRKY6* suppresses *OsPHO1;2* expression, which was further validated by enhanced *OsPHO1;2* expression in roots and improved shoot Pi accumulation in *oswrky6* knockout lines (Figure 1, 2). All these findings confirmed the role of OsWRKY6 in negatively regulating the *OsPHO1;2* expression. However, these results contradict the reported role of OsWRKY6 as an activator for *OsPR10a* and *OsPR1* genes for defence-related processes (Choi *et al*., 2015). Hence, these observations point to a dual role of OsWRKY6 in activating *OsPR10a* and suppressing *OsPHO1;2* expressions by binding to the *W-box* present in their promoters. A similar dual function has also been reported for AtWRKY42, which acts as an activator and suppressor for *AtPHT1;1* and *AtPHO1* expression, respectively (Chen *et al*., 2009; Su *et al*., 2015).

Agronomically essential cereal crops like rice, wheat, and barley have very low Pi fertilizer use efficiency (∼10-20%), leading to a bulk of fertilizers getting precipitated and drained into water bodies, causing eutrophication (Ojeda-Rivera *et al*., 2022). Thus, improving P acquisition efficiency (PAE) from soil benefits plants and the environment. The PHT1 transporters expressed in roots mainly execute Pi uptake in plants (Mudge et al., 2002; Nussaume et al., 2011), and enhancing root-associated PHTs activity gives the advantage of maximizing the utilization of Pi available in the soil. Ma *et al*., (2021) highlight that *OsPHO1;2* overexpression improves plant yield; however, they did not discuss the effect on *PHT1* expression and root morphology. Here, along with increased root-to-shoot Pi transport, *OsPHO1;2:PE* lines show improved root P content (Figure 4). We further revealed that the increase in root P is through the induction of *OsPHT1s* expression in *OsPHO1;2:PE* roots, causing more Pi uptake from media (Figure 4). Additionally, root architectural modifications such as higher root number and reduced root growth were observed in *OsPHO1;2:PE* lines (Figure 4). This is in agreement with the previous study where Pi accumulation in roots of *OsPHT1;8* overexpressing plants resulted in reduced root growth (Lapis-Gaza *et al*., 2014). We believe that reduced root growth with higher P accumulation adapted by *OsPHO1;2:PE* lines could be a trade-off mechanism of maintaining a retarded root growth to divert the carbon flow towards betterment of aerial growth. Additionally, carrying SPX and EXS domain, PHO1 acts as a Pi sensing protein along with a transporter activity (Wang *et al*., 2021). Hence, we propose that *OsPHO1;2*-driven increased Pi transport from root to shoot creates a positive driving force in roots to obtain more Pi from the external environment, thus activating root-associated *PHT1*s and increased Pi uptake. However, further research is needed to fully understand how root-associated genes, or *OsPHTs*, are directly regulated by *OsPHO1;2*.

Both *oswrky6* mutants and *OsPHO1;2:PE* lines show enhanced *OsPHO1;2* expression and improved shoot Pi accumulation compared to WT (Figure 2 and 3). However, the effect of increased *OsPHO1;2* expression on the growth pattern was not evident in *oswrky*6 mutants compared to *OsPHO1;2:PE* lines (Figure S4). Our findings offer the following explanations: (i) A TF could have multiple target genes. For example, AtWRKY42 targets *AtPHO1* and *AtPHT1;1* (Su *et al*., 2015). Hence, OsWRKY6 may act as a regulator of multiple targets related to stress and development (Choi *et al*., 2015). As a result, knocking out *OsWRKY6* could only restore average plant growth even with high P content, and (ii) multiple TFs may target the same gene (Dergilev *et al*., 2021). For example, AtWRKY6 and AtWRKY42 both suppress *AtPHO1* expression. Hence, multiple WRKY inhibitors may have a preference for the *OsPHO1;2 W-box* site in addition to OsWRKY6, and inhibitor activity of these TFs is preventing the improved growth in *oswrky6* mutants. Therefore, our genome editing approach in deleting *cis*-regulatory elements from *OsPHO1;2* promoter diminishes the chances of multiple repressors binding to *W-box* and quantitatively adjusts gene expression with enhanced grain yield.

In plants, mainly the gene promoter sequences define gene expression and modulate different adaptive developmental plasticity (Brázda *et al*., 2021). For instance, variations in the *OsSGR* promoter sequence in different *indica* rice subspecies give different senescence patterns (Shin *et al*., 2020). Also, natural variations in *ZmICE1* promoter show different behaviour toward cold treatment (Jiang *et al*., 2022). In addition to using gene editing for unravelling the function of a gene and creating desired trait, promoter editing is emerging as a powerful tool in different crop species for improving qualitative traits, increased stress tolerance, and higher yield (Shi *et al*., 2023; Tang and Zhang, 2023). A recent study by Wang *et al*. (2024) showed that deleting the repressor motif in the promoter of *NF-YC4* gene increases *NF-YC4* expression and improves protein content. In our study, we adopted a strategy of modifying promoter using a multiple gRNA approach to create a small deletion in the promoter of *OsPHO1;*2. We showed that the *OsPHO1;2:PE* lines have enhanced *OsPHO1;2* expression within roots, along with higher shoot phosphate accumulation and improved plant growth (Figure 3). Results also indicate that *OsPHO1;2:PE* lines increase yield-related traits like panicle number and overall yield without compromising grain quality (Figure 5 and 6). Such examples of improved grain yield were reported in targeting the promoter of GS3 and D18 using CRISPR/Cas12a to increase grain size and panicle number, respectively (Shi *et al*., 2023; Zhou *et al*., 2023). Our strategy is superior to earlier methods of overexpressing Pi transporters that bring Pi toxicity and growth retardation along with increasing P in plants (Ai *et al*., 2009; Wang *et al*., 2014). Therefore, we demonstrated our method of promoter editing in Pi transporter gene improves plant performance in soils treated with normal and limited P fertilizer (Figure 5, S10 and S11).

In summary, phosphate deficiency continues to be a critical factor affecting crop productivity. Different potential strategies could be exploited to improve P acquisition and utilization for crop improvement. For instance, overexpression of *PSTOL1* and phosphate transporters can increase P uptake and low P tolerance (Gamuyao *et al*., 2012; Zhang *et al*., 2014). Also, bioengineered P-solubilizing bacteria have the potential to release Pi from organic substances. Transgenic plants expressing bacterial gene *ptxD* (phosphite oxidoreductase) could metabolize phosphite (López-Arredondo and Herrera-Estrella, 2012). However, these approaches introduce transgene, which is not socially acceptable for crop plants. Our finding of CRISPR/Cas9-based promoter modification illustrated a way forward approach for editing transcription repressor sites in the promoter of *OsPHO1;2* for improving Pi transport and yield (Figure 7). The transgene-free promoter-edited plants could represent a valuable resource for commercializing rice varieties with better yield under low P input soils.

**Figure 7.**
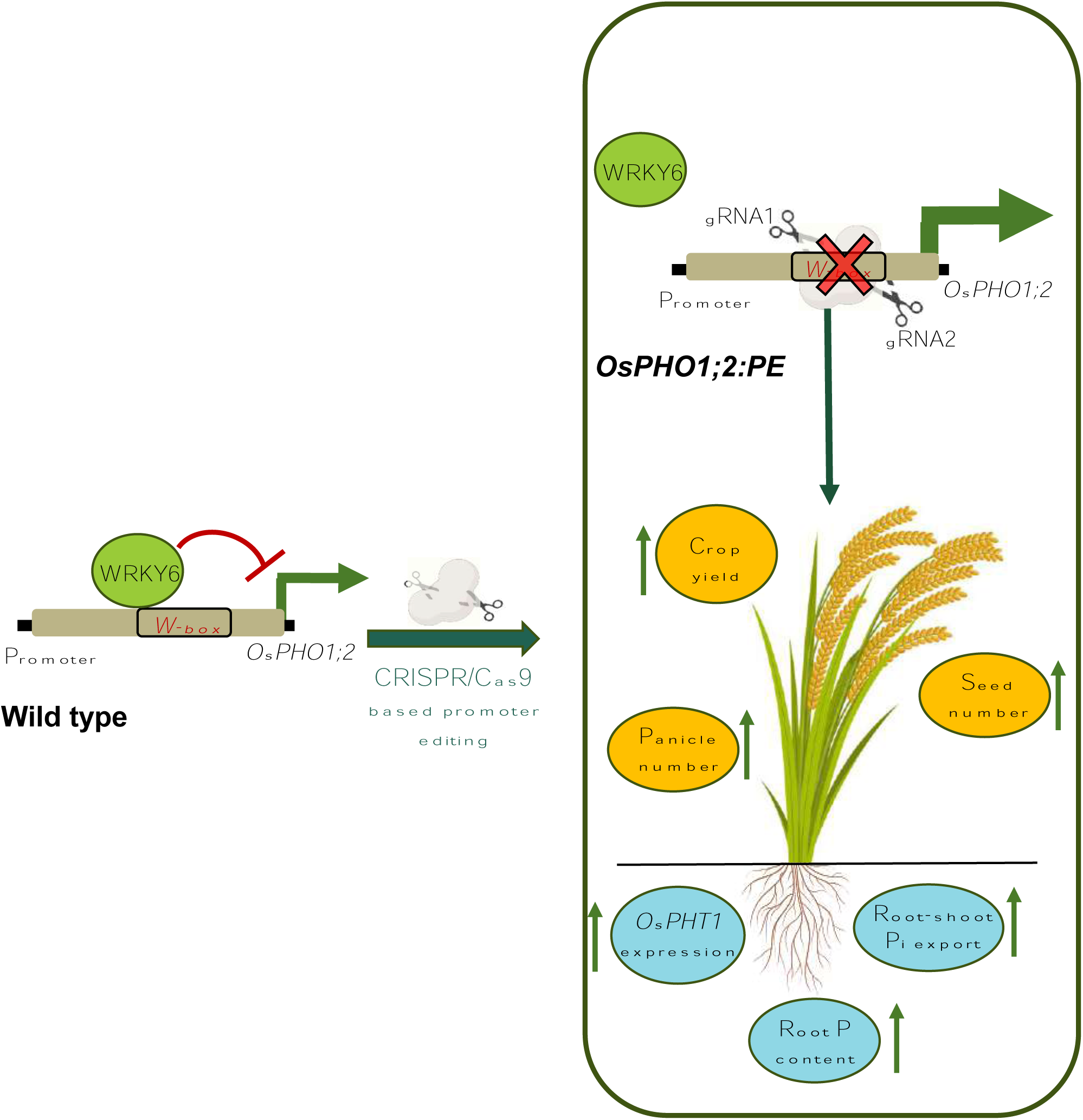
A proposed working model. In the WT rice, OsWRKY6 acts as a transcription repressor of *OsPHO1;2* by binding at the *W-box* site present in the *OsPHO1;2* promoter. Removal of *W-box* using CRISPR/Cas9 (*OsPHO1;2:PE)* alleviates OsWRKY6 and other potential inhibitors mediated inhibition, thereby increasing *OsPHO1;2* expression. Such an increase in *OsPHO1;2* transcript causes more root-shoot Pi transport, high root-associated phosphate transport (*OsPHTs*) transcript level, enhanced root Pi uptake, culminating in an overall improved crop yield.

### Experimental procedures

Dehusked rice (*Oryza sativa L. ssp. japonica*) cultivar Nipponbare (WT) seeds were surface sterilized with 0.1% HgCl_2_ and pre-germinated on half-strength Murashig and Skoog (MS) media. After four days, germinated seedlings were transferred to hydroponics Yoshida media supplemented with low P (10 μM NaH_2_PO_4_, LP) and normal P (320 μM NaH_2_PO_4_, NP) concentrations as described (Singh et al., 2015; Mehra et al., 2019). The growth assessment experiments were conducted for 21 days in a growth chamber with a day/night temperature of 30 °C/24 °C, 16 h (light)/ 8 h (dark) photoperiod (400-450 µmol photons m^−2^ s^−1^), and a relative humidity of 60%.

For gene expression analysis using RT-qPCR, seeds were inoculated in 1/4^th^ MS media in hydroponics for 9 days. Root tissues were harvested and snap-frozen in liquid nitrogen. Samples were stored at –80 °C until RNA extraction.

To investigate the seed germination percentage, surface sterilized seeds were allowed to germinate on water-soaked germination paper. The percentage was calculated at intervals of 1-4 days.

### gRNA designing, vector construction and rice transformation

For raising promoter-GUS reporter plants of *OsWRKY6*, the putative promoter (2000 bp upstream from ATG) was amplified from WT cultivar genomic DNA. The promoter amplicon was cloned upstream of the *GUS (β-glucuronidase)* reporter gene in a binary vector pMDC163 using LR*-*based cloning. The final construct was mobilized to *Agrobacterium tumefaciens* strain (EHA105). Transgenic rice plants were generated by *Agrobacterium-*mediated transformation in scutella-derived calli following the protocol described in Bhadouria *et al*., 2023. The transgenic plants were then grown in a greenhouse under the conditions mentioned above.

To generate *OsWRKY6* knockout lines, a gRNA specific to the first exon of *OsWRKY6* was designed using an online tool CRISPR GE (http://skl.scau.edu.cn/). The designed gRNA was cloned in the destination binary vector, pRGEB32 (Xie *et al*., 2015). The final construct was used to raise transgenics as described above. To identify CRISPR/Cas9 mutants, genomic DNA was extracted from the leaves using the Cetyltrimethylammonium bromide (CTAB) method (Murray and Thompson, 1980), and PCR amplification was performed using gene-specific primers followed by sanger sequencing. Positive lines were selected using Synthego-ICE tool. The selected T2 homozygous lines were used for experimentation.

For targeting *W-box* sequence (TTGACC) present in the *OsPHO1;2* promoter, two gRNAs were designed against the nearest PAM-containing sequence spanning the *W-box* region. tRNA-gRNA complex was assembled as described (Xie *et al*., 2015). Finally, the multiplex system was cloned into a binary vector, pRGEB32. The final construct was used to raise transgenics as described above. Biallelic edited plants were then selected in T2 generation by performing fragment-specific PCR followed by DNA sequencing. Transgene-free plants for downstream experiments were then selected after performing T-DNA PCR using Cas9-specific primers (Verma *et al*., 2022). All the primers used for gRNA designing and transgenic screening are detailed in Table S4.

### Analysis of off-target mutations

The CRISPR-GE tool was used to identify potential off-targets for the gRNAs used to generate *OsPHO1;2:PE* lines. Primers were designed to amplify potential off-target regions. The amplified PCR product was purified and directly sent for Sanger sequencing. Off-target mutations were identified using Synthego tool (Synthego, 2019). The promoter-edited lines with non-detectable off-target mutation were selected for further use. Primers used for screening are mentioned in Table S4.

### Gene expression analysis

Total RNA from plant tissue was extracted using Trizol reagent (Invitrogen, USA). 1 μg RNA was reverse transcribed into cDNA using a high-capacity Reverse transcription kit (Applied biosystems, USA). The synthesized cDNA was then used to perform real-time quantitative PCR (RT-qPCR) using SYBR® Green (Applied Biosystems, USA) master mix in Applied Biosystems 7500 Fast Real-Time PCR instrument. Rice housekeeping gene *Ubiquitin5* was used as an endogenous control. The relative gene expression level was calculated using ΔΔCt method. Primers used in RT-qPCR are shown in Table S4.

### Soluble Pi and total P content measurement

Soluble Pi estimation of fresh tissue was performed as described (Chiou *et al*., 2006). Briefly, samples were crushed in liquid nitrogen and homogenized in extraction buffer (10 mm Tris– HCl pH 8.0, 1 mm EDTA, 0.1 M NaCl, 1 mM β marcaptoethanol) in 1:10 ratio. The solution was diluted five times in CH_3_COOH followed by 30 minutes of incubation at 42 °C. Next, 300 μl of solution was mixed with 700 μl of assay solution (10% Ascorbic acid, 0.42% ammonium molybdate in 2N H_2_SO_4_) with 30 minutes of incubation at 42 °C. The absorbance was then recorded at 820 nm wavelength using POLAR star Omega plate reader (BMG Labtech, Germany). Pi concentration was measured using a standard solution of 50 ppm KH_2_PO_4._

The yellow vanadomolybdate method was followed to measure total P in root and shoot tissues (Mehra et al. 2017). Briefly, tissues were oven-dried completely at 60 °C for three days and flamed to ash at 500 °C overnight. Ash was dissolved in 2N HCl and mixed with an ammonium metavanadate solution to obtain a yellow-coloured complex. Absorbance was estimated at 410 nm wavelength in a microplate reader using 50 ppm KH_2_PO_4_ solution as a standard.

For elemental analysis (P, Fe) in seeds, brown rice powder (200 mg) was digested with 8ml of 65% HNO_3_ at 180 °C in a microwave digestion system. The samples were diluted, filtered with 0.2 μm filters, and measured with an Inductively coupled plasma mass spectrometry (ICP-MS), Agilent 7800.

### Measurement of ^33^P uptake and transport

For measuring ^33^P transfer from root to shoot, dehusked sterilized seeds of WT and *OsPHO1;2:PE* lines were germinated for nine days in 1/4^th^ of MS hydroponics solution. Roots of seedlings were washed with distilled water and immersed in ^33^P radioactive solution (3.1 μCi/mL; 10 μM Pi) for 2 hours. After incubation, roots were washed with 10 mM cold phosphate solution until all external radioactivity was removed. Radioactive phosphate was separately measured for root and shoot in a scintillation counter. P transfer was then calculated by dividing shoot radioactivity by total radioactivity in plants. To obtain ^33^P uptake kinetics within the plant, the total plant radioactivity per mg fresh weight of roots was calculated.

### Root system architecture traits quantification

Seeds were pregerminated on 1/4^th^ MS media for four days and transferred to hydroponics in Yoshida media supplemented with low P (10 μM NaH_2_PO_4_, LP) and normal P (320 μM NaH_2_PO_4_, NP) concentrations. Roots were harvested after 21 days of treatment and fixed in FAA fixative solution (10% formaldehyde, 50% ethanol, and 5% acetic acid). For measurement, roots were stained with neutral red dye (0.5 mg/ml), followed by scanning using EPSON Perfection V850 Pro scanner in the professional mode. Finally, scanned images were analysed using RhizoVision Explorer v2.0.3 (Seethepalli and York, 2021) by applying prescribed algorithms (Seethepalli et al., 2021).

### Electrochemical mobility shift assay (EMSA)

The DNA binding domain of OsWRKY6 (222-380 amino acid) was amplified using cDNA from rice seedlings and cloned in pMALc2x vector for fusion with N terminal MBP tag. The WRKY-MBP were expressed in *E. coli* strain BL21 with the induction of 0.2mM IPTG for 12-14 hrs at 22 °C and further purification with MBP resin. The oligo duplex (30 bp) containing the WRKY binding site *W-box* element (‘TTGACC’) and the mutant probe (‘TTGACC’ was replaced by ‘AAAAAA’) was labelled using DIG Oligonucleotide 3ʹ-End Labelling Kit (Roche, Germany). The 20 ng of labelled probes were incubated for 30 minutes at 22 °C with 5 μg recombinant protein with binding buffer (1 μg poly(dI-dC), 15 mM HEPES (pH 8), 1 mM MgCl2, 30 mM KCl, 0.02 mM DTT, 0.2 mM EDTA, and 0.6% glycerol). The reaction solution was run on native polyacrylamide gel (4% v/v) for 3 hours and then transferred to a positively charged nylon membrane, followed by UV crosslinking. Band shifts were finally detected on X-ray film using Anti-Digoxigenin-AP, Fab fragments kit (Roche). Primers and probes used in the experiment are listed in Table S4.

### Subcellular localization and histochemical GUS staining

The full-length coding sequence of OsWRKY6 with the stop codon was amplified and cloned into the pSITE3cA vector to generate 35S:eYFP-OsWRKY6. This construct and the nuclear marker 35S:RFP-NLS were co-transformed into *N. benthamiana* leaves by *A. tumefaciens* (EHA 105) mediated infiltration method as described (Zhang *et al*., 2020). The fluorescence signal was observed under a confocal laser scanning microscope (Leica TCS SP5, Leica microsystem, Germany). Primers used in the experiment are listed in Table S4.

Histochemical localization of GUS activity in *OsWRKY6p*:*GUS* plants was performed as described earlier (Jefferson et al., 1987). Briefly, different tissues from *OsWRKY6p*:GUS transgenics were incubated in GUS staining buffer containing the GUS substrate (1 mm 5-bromo-4-chloro-3-indolyl-β-d-glucuronide) at 37 °C for 2-3 days. Following the development of colour, tissues were immersed overnight in 70%(v/v) ethanol for the removal of chlorophyll. Images of tissues were captured under the Leica S9i light microscope.

### Transient expression assay

The transient expression assay was carried out in rice seedlings as described earlier (Singh *et al*., 2023). Briefly, 1.5 kb putative promoter region of *OsPHO1;2* which includes the *W-box*, was cloned in the pCAMBIA1303 vector by replacing CaMV35S promoter upstream of GUS gene and transformed into *Agrobacterium* (EHA 105). The cells were grown overnight at 28°C. When the OD reached 0.6-0.8, the cells were harvested and resuspended in infiltration buffer (10 mM MES, 10 mM MgCl_2_, and 100 mM acetosyringone). The 3-day-old rice seedlings (WT and *oswrky6)* were incubated in *Agrobacterium* culture and then vacuum infiltrated. The seedlings were kept in a growth chamber for 2-3 days, and then GUS activity was analysed. GUS staining of roots was performed as described above. Primers used in the experiment are listed in Table S4.

For quantification of GUS activity, the fluorometric MUG assay was performed as described (Francis and Spiker, 2005). Briefly, the total protein from rice roots was isolated using GUS extraction buffer (150 mM sodium phosphate at pH 7.0, 10 mM EDTA, 10 mM β-mercaptoethanol, 0.1% Triton X-100, 140 μM PMSF and 0.1% sarcosyl). 13 µg of isolated protein from WT and *oswrky6* roots was incubated with GUS assay buffer (GUS extraction buffer containing 1.2 mM 4-methylumbelliferyl β-D-glucuronide (MUG), Sigma) at 37 °C in the dark. After 1 hour of incubation, 10 µL of the reaction mix was added to 190 µL of stop buffer (200 mM sodium carbonate) in a 96-well opaque microplate. 4-MU (4-methylumbeliferone) was used as a standard. The fluorescence was measured in POLARstar Omega plate reader with 355 nm (excitation) and 450 nm (emission). GUS activity was mentioned as nM/min/mg protein.

### Starch and amylose estimation

Dehusked seeds were dried and powdered to estimate the total starch in rice seeds. Starch estimation was performed according to the Total Starch (AA/AMG) Assay Kit (Megazyme, Ireland) and all steps were followed as per manufacturer’s instructions. The available total starch was calculated using the Mega-Calc™ software tool.

The iodine: potassium iodide (I_2_:KI) procedure was used to estimate amylose (Juliano, 1971). Briefly, 100 mg of crushed seeds were mixed with 20 μl of 95% ethanol and 180 μl of 1M NaOH and incubated in boiling water for 10 minutes. The volume was made up to 2 ml. The 100 μl solution was mixed with 20 μl acetic acid and 40 μl 0.2% I_2_-KI reagent. Absorbance was then recorded at 620 nm using POLAR star Omega plate reader (BMG Labtech, Germany). The amylose content was calculated using a standard graph (Avaro *et al*., 2011).

For observing the morphology of starch granules, dehulled seeds were crushed into a fine powder using a tissue lyser followed by staining with Lugol solution (iodine: potassium iodine solution diluted in deionized water [1:10]; Himedia) for at least 10 seconds. The starch granules were then examined under a light microscope.

The packaging of starch granules was visualized using a scanning electron microscope (SEM, EVO^®^ 911 LS10 Zeiss, Germany). Matured seeds were cut from the center of the endosperm and photographs were taken at magnification of 2500x.

### Phytic acid measurement

Phytic acid was measured in rice seeds using Phytic acid assay kit (Megazyme K-PHYT, Ireland) following the manufacturer’s instructions. Briefly, 200 mg of crushed seeds were taken in glass tubes and 2 ml of 0.66 M HCl was added. The tubes were then incubated at 28 °C shaker at 200 rpm for 5-6 hours. 1ml of extraction was centrifuged at 13000 rpm and supernatant (0.5 ml) were then taken in another tube with a subsequent addition of 0.75 M NaOH (0.5 ml) to neutralize it. The following steps to measure free and total Phosphorus were followed as described (Megazyme K-PHYT, Ireland). Calculation for phytic acid was done using the Mega-Calc™ software tool (Megazyme K-PHYT).

## Statistical Analysis

All numerical data are presented as means with a standard error bar. Statistical significance was determined by pairwise Student’s *t*-test using the Microsoft Office Excel program. Statistical significance was indicated: * for P ≤ 0.05, ** for P ≤ 0.01, and *** for P ≤ 0.001. Data from Figure 1a was analysed by Brown-Forsythe and Welch’s one-way ANOVA test, followed by the multiple comparison with P value (P < 0.05) adjusted with FDR.

## Author contributions

K.M. conducted most experiments, analyzed data, and wrote the manuscript. B.M. conducted experiments, analyzed data and edited the manuscript. B.S., U.S. and A.J. helped with the experiments. S.S., R.P., H.V.T. and S.K.M. helped with soil-based experiments. J.G. and Y.P. conceived the project, J.G. supervised experiments, analyzed data, and finalized the manuscript with help from Y.P.

## Acknowledgments

This research is funded by the Indo-Swiss joint research project (BT/IN/Swiss/46/JG/2018-2019 to JG and IZLIZ3_183114/1 to YP) and a grant from the Herbette Foundation of the University of Lausanne. Research fellowship to K.M. from CSIR, India; B.M. from DBT, India and B.S. and U.S from NIPGR, India are gratefully acknowledged.

## Data availability statement

All relevant data generated or analysed are included in the manuscript with supporting materials.

## Conflict of interest

The authors declare no conflicts of interest.

## Supplementary figures legends

Figure S1. Subcellular location of OsWRKY6 in *N. benthamiana* leaves.

Figure S2. Generation of os*wrky6* knockout mutants using CRISPR/Cas9.

Figure S3. *OsWRKY6* expression levels in *oswrky6* knockout mutants.

Figure S4. Phenotypic analysis of os*wrky6* knockouts.

Figure S5. Genotypic analysis of *OsPHO1;2:PE* plants.

Figure S6. Cas9 detection in T2 generation of *OsPHO1;2:PE* marker-free lines.

Figure S7. Total shoot P concentration and shoot dry biomass in WT and *OsPHO1;2:PE* lines.

Figure S8. Total root P concentration and root efficiency in WT and *OsPHO1;2:PE* lines.

Figure S9. Evaluation of agronomic traits in WT and *OsPHO1;2:PE* lines (Year 2023).

Figure S10. Evaluation of agronomic traits in WT and *OsPHO1;2:PE* lines (Year 2024).

Figure S11. Evaluation of agronomic traits in WT and *OsPHO1;2:PE* lines under low P fertilizer supply.

Figure S12. Amylose content quantification and starch granule visualization in seeds of WT and *OsPHO1;2:PE* lines.

Figure S13. Evaluation of grain physiological P use efficiency (PPUE) in WT and *OsPHO1;2:PE* lines.

Figure S14. *OsPHO1;2:PE* lines show no alteration in seed Fe content.

## Supporting tables

Table S1. Genes co-expressing with *OsPHO1;2*.

Table S2. Putative CRISPR/Cas9 off-targets for gRNAs used in the generation of *OSPHO1;2:PE* lines.

Table S3. Off-targets (OT) prediction for gRNAs used in the generation of *OsPHO1;2:PE* lines. Table S4. List of the primers and probes used in this study.

## Notes

### Competing Interest Statement

The authors have declared no competing interest.

