## Supplementary figures for "Editing *cis*-elements of *OsPHO1;2* improved phosphate transport and yield in rice"

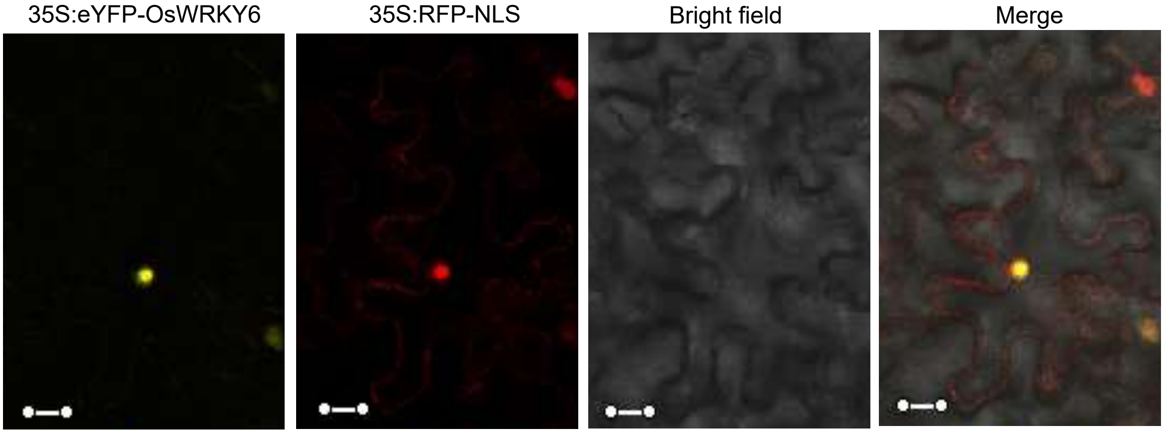


Figure S1. **Subcellular localization of OsWRKY6 in *N. benthamiana* leaves**. Transient co-expression of 35S:eYFP-OsWRKY6 and 35S:RFP-NLS in *N. benthamiana* leaves. 35S:RFP-NLS was used as a nuclear localized marker. Scale bar: 20 μm.

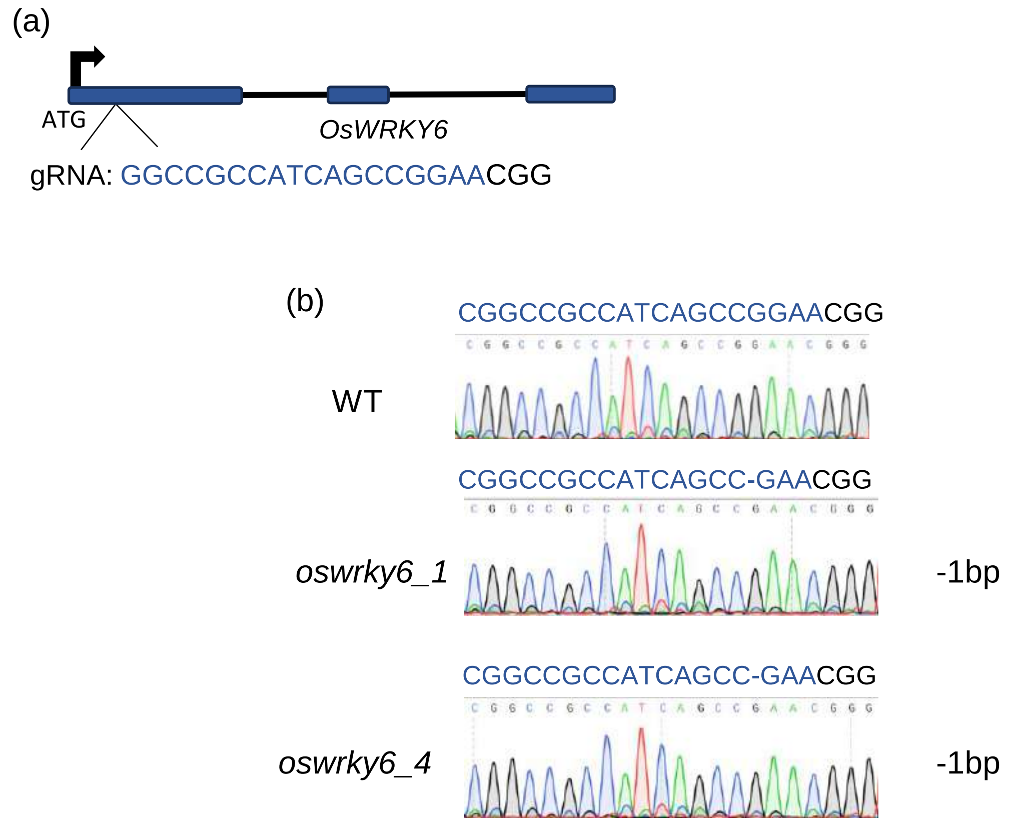


Figure S2. **Generation of os*wrky6* knockout mutants using CRISPR/Cas9**. (a) Schematic representation of *OsWRKY6* gene structure comprising three exons and two introns. Exons are represented by blue bars separated by black lines representing introns. The arrow represents the translation initiation codon (ATG). gRNA was designed from the first exon. (b) Sanger-sequencing chromatographs are shown for selected T2 generation *oswrky6* knockout with biallelic mutation. Sanger-sequencing was performed directly for the amplified product containing gRNA region. gRNA sequence is highlighted in blue colour. The three nucleotides in black color indicate PAM sequence. The WT gene sequence is shown for reference, and deletions in sequences of both alleles are labeled as "–".

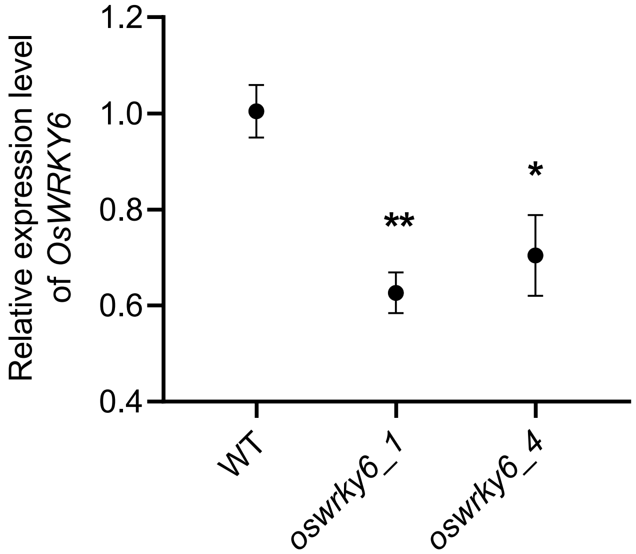


Figure S3. ***OsWRKY6* expression levels in *oswrky6* knockout mutants**. Expression level of *OsWRKY6* in root tissue of 9 days-old seedlings of wild type (WT) and *oswrky6* knockout lines. Relative expression was calculated relative to WT. *Ubiquitin5* was used as an endogenous control. Data represent means ± SE (n= 4, each replicate contains a pool of 5 seedlings). The Student's *t*-test determined significant changes. * and ** indicate significant differences from control (WT) at P-value ≤0.05 and ≤0.01, respectively.

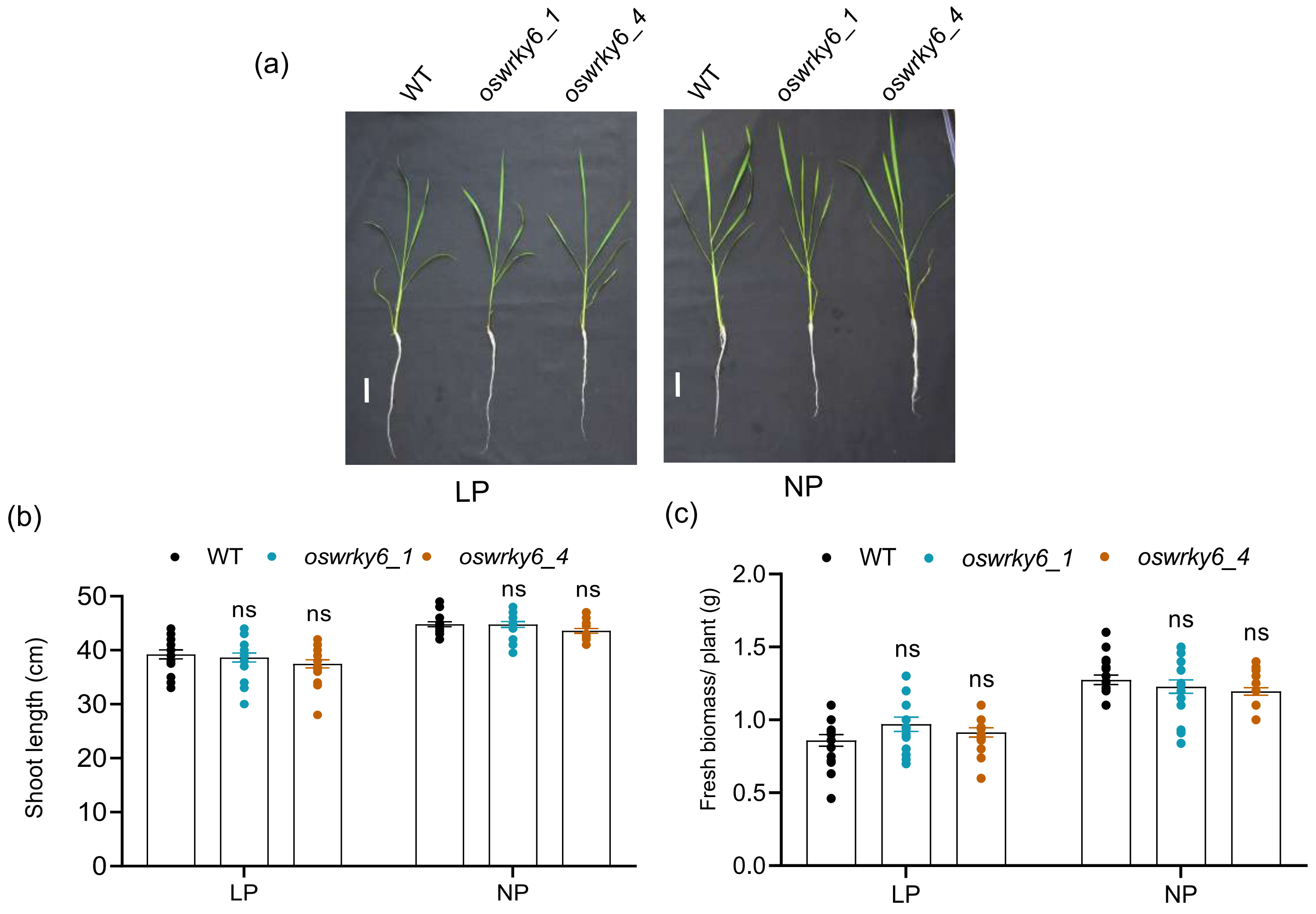


Figure S4. **Phenotypic analysis of *oswrky6* knockouts**. (a) Phenotypic growth analysis, (b) shoot length and (c) plant biomass of 21 days-old seedlings of wild type (WT) and *oswrky6* lines grown in Yoshida media with low (10 µM $NaH_2PO_4$, LP) and normal P (320 µM $NaH_2PO_4$, NP) conditions. Data represent means ± SE (n=15-20). Each dot represents one biological replicate. Significant changes were determined by the Student's *t*-test. ns indicate no significant difference.

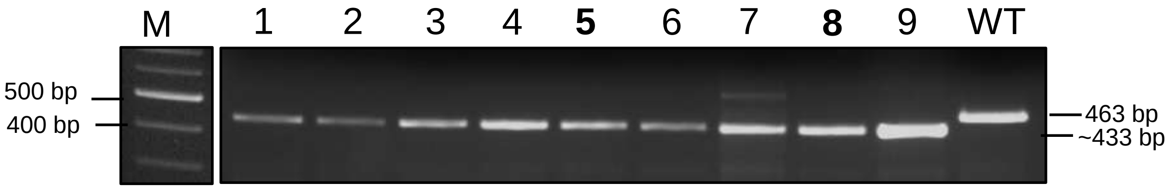


Figure S5. **Genotypic analysis of *OsPHO1;2:PE* lines**. Genomic DNA PCR of progenies from T2 generation of *OsPHO1;2:PE* lines using primers spanning the *W-box* region. The expected band size for WT and *OsPHO1;2:PE* lines are 463 bp and ~433 bp, respectively. Each line number is shown on the top of the gel, and the selected line for further experimentation is denoted in bold. M- DNA ladder; bp- base pairs, WT- wild type.

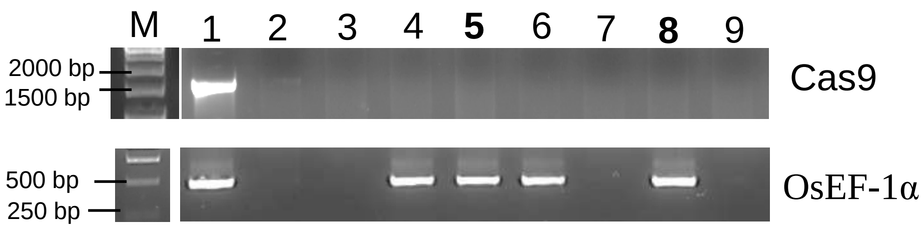


Figure S6. **Cas9 detection in T2 generation of *OsPHO1;2:PE* marker-free lines.** Biallelic *OsPHO1;2:PE* lines of T2 generation were screened for transgene-free nature by employing genomic-DNA PCR using Cas9 primers. House-keeping gene elongation factor 1 alpha (*EF-1α*) PCR was performed to assess the quality of genomic DNA. Each line number (T2 generation) is shown on the top of the gel, and the selected line for further experimentation is denoted in bold. M- DNA ladder; bp- base pairs.

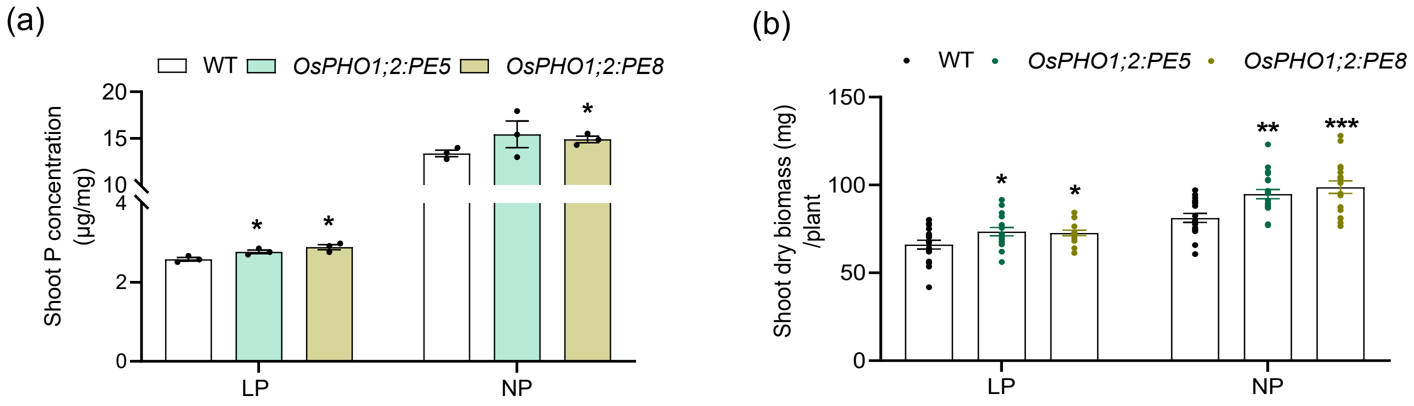


Figure S7. **Total shoot P concentration and shoot dry biomass in WT and *OsPHO1;2:PE* lines.** (a) Total P (n= 3, each replicate contains a pool of 5 seedlings) and (b) shoot dry biomass (n=15-20) in dried shoot tissues of 21-days-old seedlings of wild type (WT) and *OsPHO1;2:PE* lines grown in Yoshida media with low (10 µM $NaH_2PO_4$, LP) and normal P (320 µM $NaH_2PO_4$, NP) conditions. Data represent means ± SE. Each dot represents one biological replicate. Significant changes were determined by the Student's *t*-test. *, ** and *** indicate significant differences from control (WT) at P-value ≤0.05, ≤ 0.01 and ≤0.001, respectively.

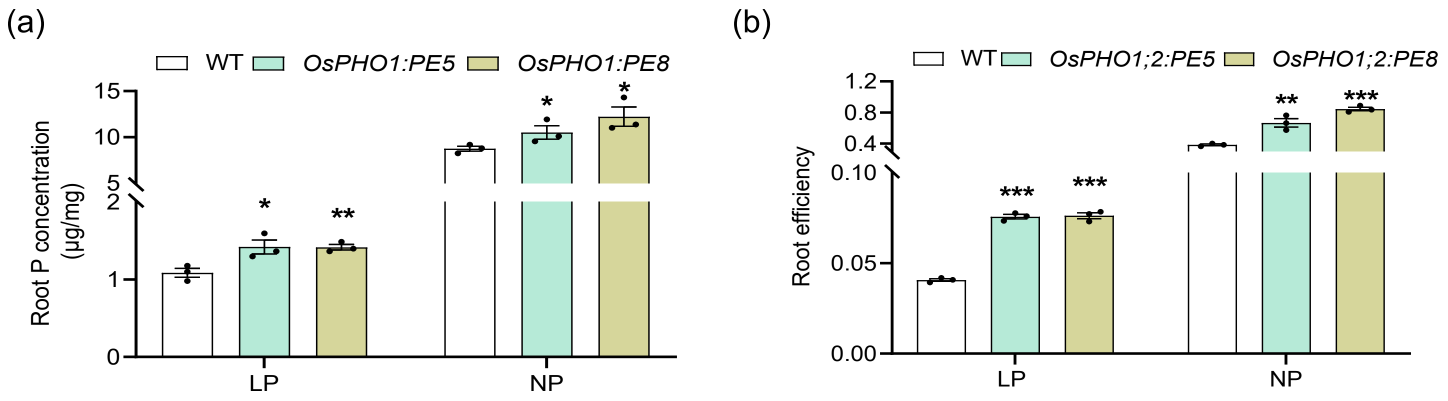


Figure S8. **Total root P concentration and root efficiency in WT and *OsPHO1;2:PE* lines.** (a) Root P concentration and (b) root efficiency measured in 21 days-old seedlings of wild type (WT) and *OsPHO1;2:PE* lines grown in Yoshida media with low (10 µM $NaH_2PO_4$, LP) and normal P (320 µM $NaH_2PO_4$, NP) conditions. Root efficiency is calculated as plant total phosphate (root P concentration x root dry weight) + (shoot P concentration x shoot dry weight) per root surface area ($mm^2$). Data represent means ± SE (n= 3, each replicate contains a pool of 5 seedlings). Each dot represents one biological replicate. Significant changes were determined by the Student's *t*-test. *, ** and *** indicate significant differences from control (WT) at P-value ≤0.05, ≤ 0.01 and ≤0.001, respectively.

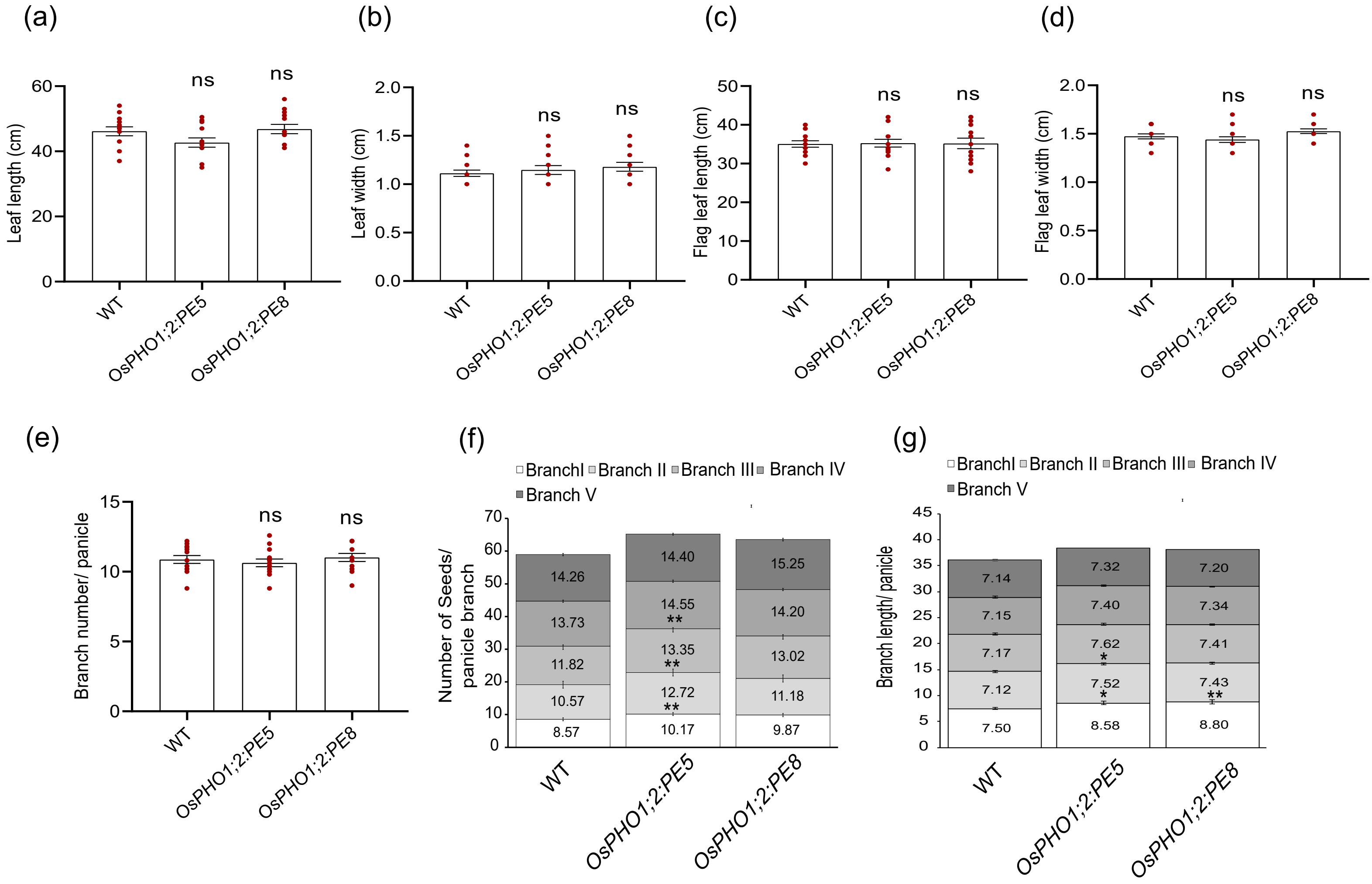


Figure S9. **Evaluation of agronomic traits in WT and *OsPHO1;2:PE* lines (Year 2023).** Measurement of (a) leaf length, (b) leaf width, (c) flag leaf length, (d) flag leaf width, (e) branch number/ panicle, (f) seed number/ panicle branch and (g) branch length/ panicle in wild type (WT) and *OsPHO1;2:PE* lines at maturity. Data represent means ± SE (n = 12-15). Each dot represents one biological replicate. Significant changes were determined by the Student's *t*-test. ns indicate no significant difference. * and ** indicate significant differences from control (WT) at P-value ≤0.05 and ≤0.01, respectively.

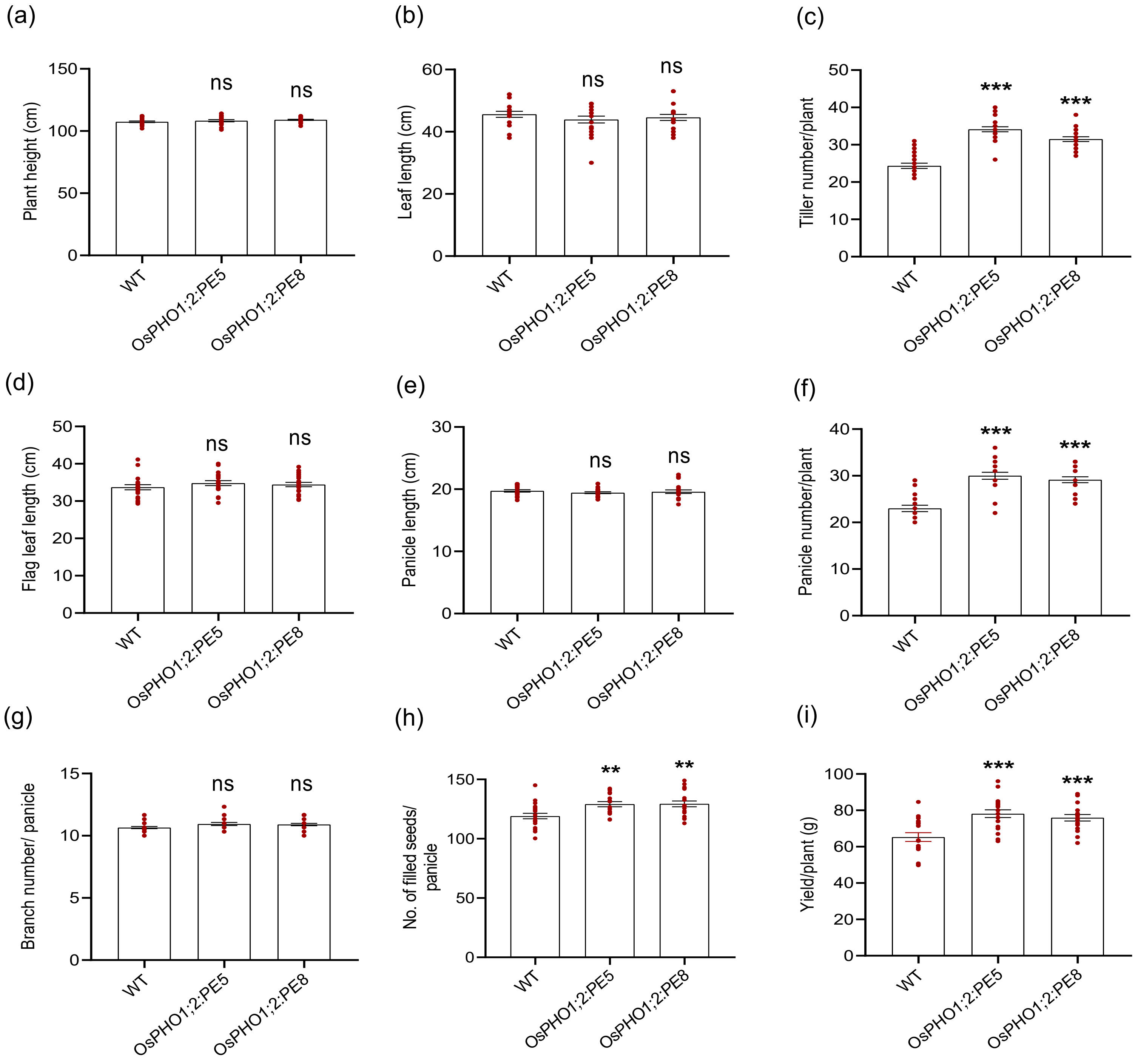


Figure S10. **Evaluation of agronomic traits in WT and *OsPHO1;2:PE* lines (Year 2024).** Measurement of (a) plant height, (b) leaf length, (c) tiller number/plant, (d) flag leaf length, (e) panicle length, (f) panicle number/plant, (g) branch number/ panicle, (h) number of filled seeds/ panicle and (i) yield/plant in wild type (WT) and *OsPHO1;2:PE* lines at maturity. Data represent means ± SE (n = 17-20). Each dot represents one biological replicate. Significant changes were determined by the Student's *t*-test. ns indicate no significant difference. ** and *** indicate significant differences from control (WT) at P-value ≤0.01 and ≤0.001, respectively.

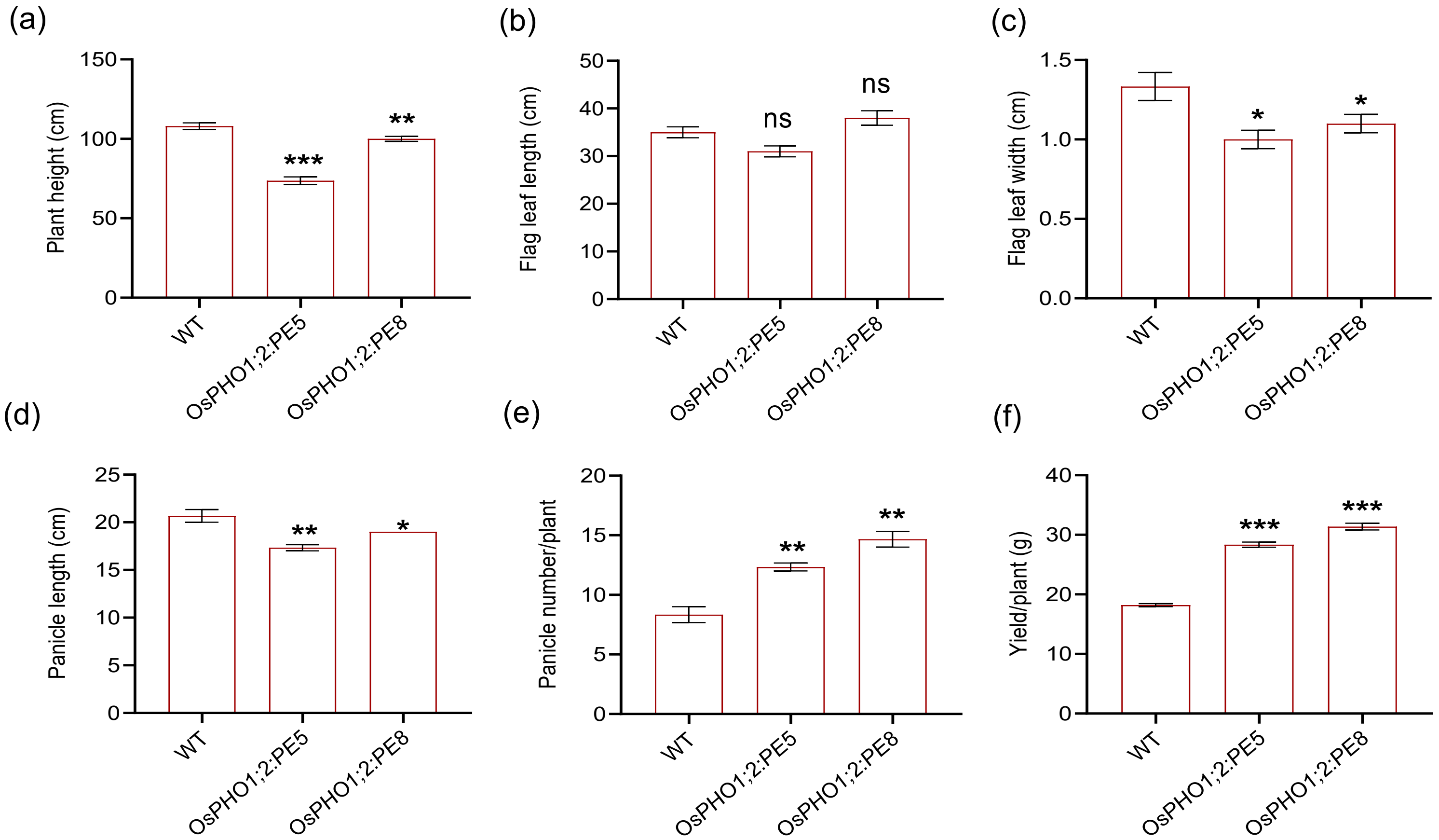


Figure S11. **Evaluation of agronomic traits in WT and *OsPHO1;2:PE* lines under low P fertilizer supply.** Measurement of (a) plant height, (b) flag leaf length, (c) flag leaf width, (d) panicle length, (e) panicle number, (f) yield/plant in wild type (WT) and *OsPHO1;2:PE* lines grown under low P soil. Data represent means ± SE (n = 3 independent replicates). Significant changes were determined by the Student's *t*-test. ns indicate no significant difference. *, ** and *** indicate significant differences from control (WT) at P-value ≤0.05, ≤0.01 and ≤0.001, respectively

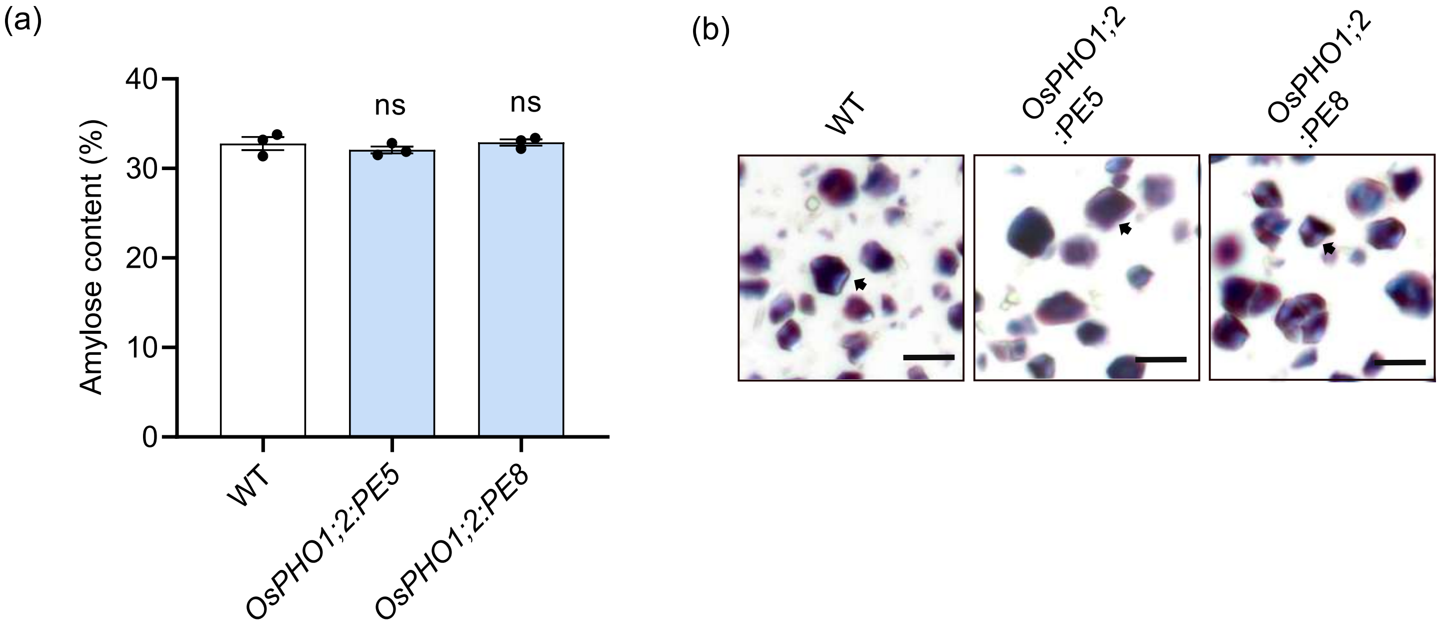


Figure S12. **Amylose content quantification and starch granule visualization in seeds of WT and *OsPHO1;2:PE* lines.** (a) Amylose content in brown seeds of wild type (WT) and *OsPHO1;2:PE* lines. Data represent means ± SE (n=3, each replicate contains a pool of 10-15 seeds). (b) Light microscopy of iodine-stained starch granules in powder of brown seeds of WT and *OsPHO1;2:PE* lines. The arrow indicates polygonal starch granules. Each dot represents one biological replicate. Significant changes were determined by the Student's *t*-test. ns indicate no significant differences from control (WT). Scale bar: 20 µm.

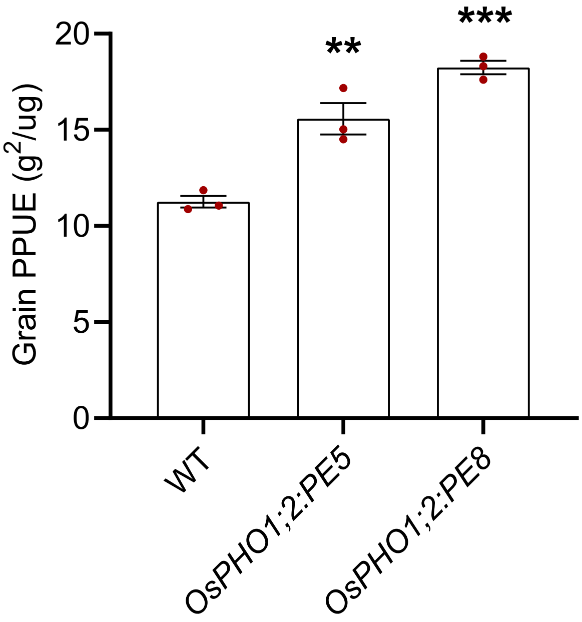


Figure S13. **Evaluation of grain physiological P use efficiency (PPUE) in WT and *OsPHO1;2:PE* lines.** Measurement of grain PPUE in wild type (WT) and *OsPHO1;2:PE* lines. Grain PPUE is calculated as yield (g) per seed P concentration (mg/g). Data represent means ± SE (n = 3). Each dot represents one biological replicate. Significant changes were determined by the Student's *t*-test. ** and *** indicate significant differences from control (WT) at P-value ≤0.01 and ≤ 0.001, respectively.

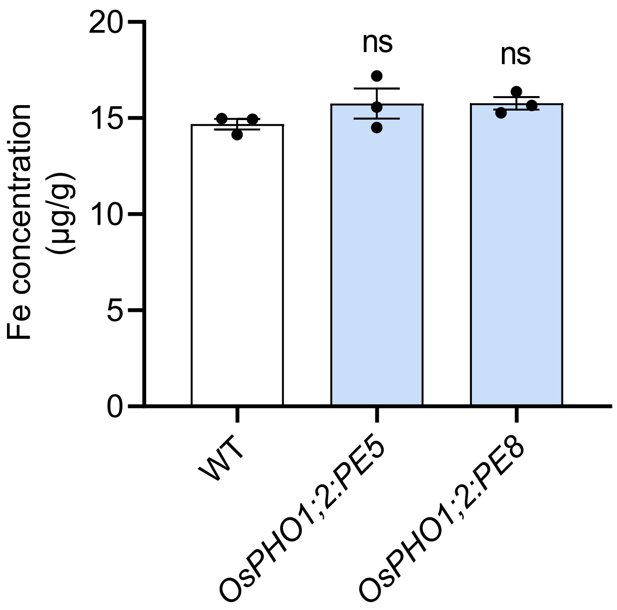


Figure S14. ***OsPHO1;2:PE* lines show no alteration in seed Fe concentration.** Fe concentration in brown seeds of wild type (WT) and *OsPHO1;2:PE* lines measured by Inductively Coupled Plasma Mass Spectrometry (ICP-MS). Data represent means ± SE (n=3, each replicate contains a pool of 10-15 seeds). Each dot represents one biological replicate. ns indicate no significant differences from control (WT). Significant changes were determined by the Student's *t*-test.
